# A chromatin-associated regulator of RNA Polymerase III assembly at tRNA genes revealed by locus-specific proteomics

**DOI:** 10.1101/2023.04.17.534528

**Authors:** Maria Elize van Breugel, Ila van Kruijsbergen, Chitvan Mittal, Cor Lieftink, Ineke Brouwer, Teun van den Brand, Roelof J.C. Kluin, Renée Menezes, Tibor van Welsem, Andrea Del Cortona, Muddassir Malik, Roderick Beijersbergen, Tineke L. Lenstra, Kevin Verstrepen, B. Franklin Pugh, Fred van Leeuwen

## Abstract

Transcription of tRNA genes by RNA Polymerase III (RNAPIII) is tightly regulated by signaling cascades in response to nutrient availability. The emerging notion of differential tRNA gene regulation implies the existence of additional regulatory mechanisms. However, tRNA gene-specific regulatory factors have not been described. For that reason, we decoded the proteome of a single native tRNA gene locus in yeast. We observed dynamic reprogramming of the core RNAPIII transcription machinery upon nutrient perturbation. In addition, we identified Fpt1, a protein of unknown function. Fpt1 uniquely occupied tRNA genes but its occupancy varied and correlated with the efficiency of RNAPIII eviction upon nutrient perturbation. Decoding the proteome of a tRNA gene in the absence of Fpt1 revealed that Fpt1 promotes eviction of RNAPIII. Cells without Fpt1 also showed impaired shutdown of ribosome biogenesis genes upon nutrient perturbation. Our findings provide support for a chromatin-associated mechanism required for RNAPIII eviction from tRNA genes and for tuning an integrated physiological response to changing metabolic demands.

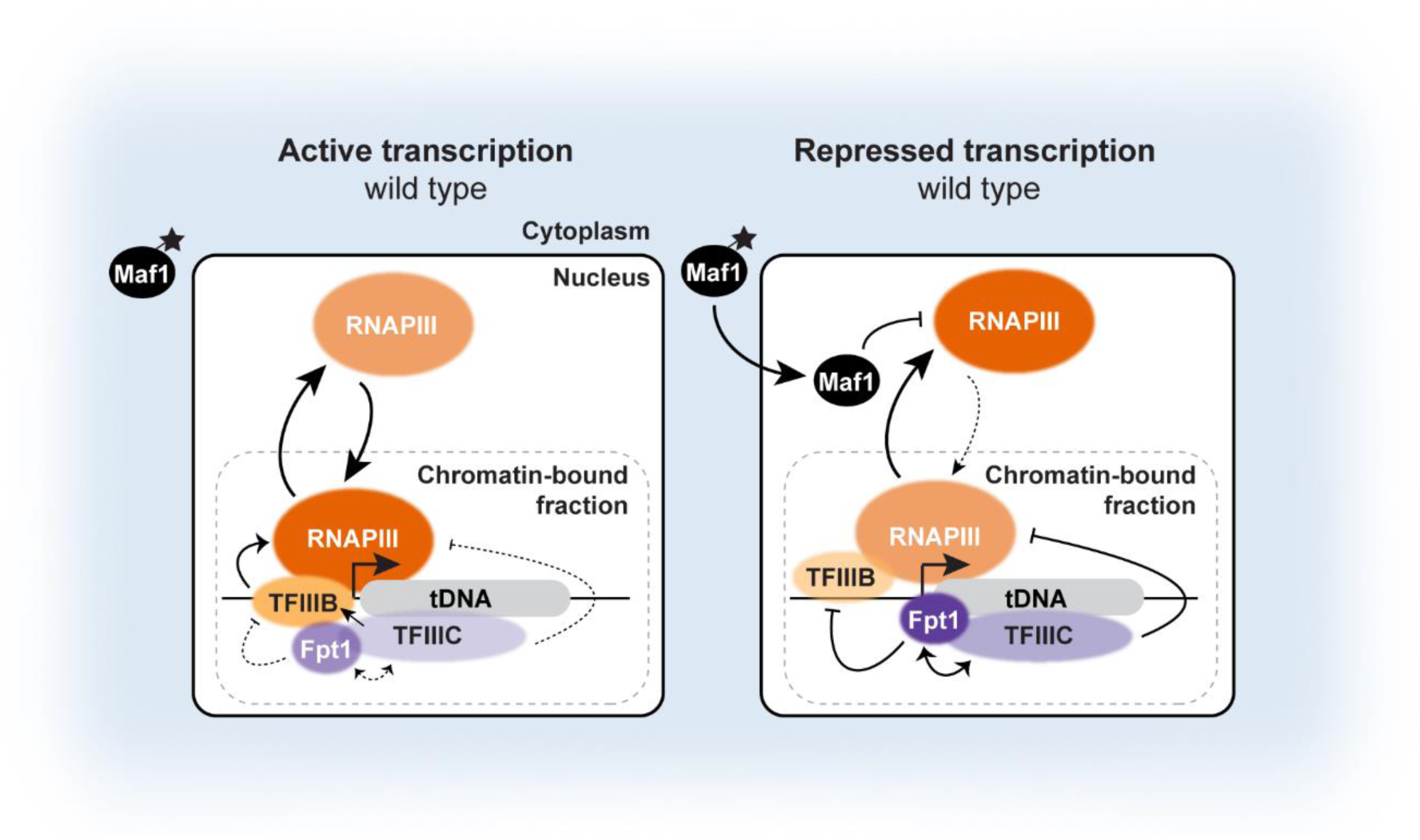

## INTRODUCTION

Eukaryotic gene transcription is a fundamental cellular process that requires tight regulation. Gene transcription is facilitated by RNA polymerase enzyme complexes that collaborate with transcription factors, repressors, chromatin remodelers, and many other cellular factors [1–3]. RNA polymerases are categorized in three classes, each specialized in transcription of a different subset of genes [4]. RNA Polymerase III (RNAPIII) mainly transcribes tDNAs, short redundant DNA fragments that code for transfer-RNAs (tRNAs) [1, 2, 5]. While tDNAs compromise a small fraction of the genome, transcription of tRNA genes accounts for ∼15% of the total RNA pool [1]. Transfer-RNA molecules function as amino acid deliverers in the process of mRNA translation but recent studies emphasize that tDNAs and the RNAs they encode are involved in a wide variety of processes such as genome organization, aging, cancer, and other diseases [6–10]. In agreement with their cellular abundance and important biological roles, tRNAs are tightly regulated at multiple levels, from transcription, post-transcriptional processing and modification, to transport and degradation [1, 5, 11–14]. Remarkably, chromatin-associated regulatory mechanisms of tDNA transcription are still poorly understood, which sharply contrasts with the wealth of knowledge on regulation of RNA polymerase II (RNAPII) [2, 3, 5, 15].

From a core transcription perspective, tDNAs are small, self-contained elements [16–18]. Transcription of tDNAs is under the control of A- and B-box promoter elements that are located within the gene body coding for the tRNA. In nutrient-rich conditions, assembly factor TFIIIC binds the internal A- and B-box promoter elements and recruits the transcription factor TFIIIB, which promotes binding of RNAPIII to the upstream transcription start site to initiate transcription. Upon transcription initiation, elongation, and termination, RNAPIII dissociates from the tDNA or reinitiates another cycle of transcription in which the same RNAPIII molecule is recycled [5, 19]. In mammalian cells, the global activator MYC activates tDNA transcription in nutrient-rich conditions [15, 20]. In repressive conditions, such as nutrient deprivation, tDNA transcription is repressed by Maf1 [5]. Maf1 is a conserved and well-characterized repressor originally identified in *Saccharomyces cerevisiae* by a classical genetic approach [21]. In nutrient-rich conditions, Maf1 is inactivated by phosphorylation depending on upstream kinases such as TORC1 [5, 19]. Inactive Maf1 is retained in the cytoplasm but in repressive conditions, Maf1 is dephosphorylated and imported into the nucleus where it represses RNAPIII transcription *in Trans* in a two-step fashion. First, Maf1 prevents de-novo assembly of TFIIIB onto the DNA by interaction with TFIIIB. Second, Maf1 represses RNAPIII recruitment to the DNA by direct interactions with RNAPIII [22, 23]. As a consequence, upon nutrient deprivation and other stress signals, RNAPIII recruitment is blocked and tDNA transcription is halted.

Interestingly, tRNA genes at different genomic locations but with identical gene bodies, and hence identical A- and B-box promoter elements, can show different expression dynamics. This was first observed in the yeast genome where in repressive conditions (e.g. growth on a non-fermentable carbon source), expression of nearly all tRNA genes was reduced but a small subset of tRNA genes was significantly less repressed and less dependent on Maf1 for reasons still not known [24–26]. Non-homogenous transcriptional regulation of tRNA genes has also been observed in human cells (e.g. [27–31]). Indeed, the relative abundance of individual tRNAs varies considerably across tissues and cell lines, in cells engaged in proliferation or differentiation, and in pathologies such as cancer [32, 33]. These observations suggest that regulatory mechanisms outside the tDNA elements or embedded in the tDNA chromatin must be at play to tune RNAPIII activity in response to changing cellular demands [34]. However, cis-regulatory elements or transcription- and chromatin-factors dedicated to gene-specific tDNA transcription by RNAPIII have not been described [5, 15, 35, 36].

Identifying regulators of tRNA genes is hampered by many technical challenges. For example, many tRNA genes occur in multiple copies with identical body sequences across the genome, precluding mapping of mature tRNA sequences back to their tRNA gene of origin [37]. In addition, tRNA genes are prone to non-specific cross-linking of proteins, presumably due to their high level of transcription and open chromatin structure [38]. Moreover, tRNAs are highly structured and harbor many post-transcriptional base modifications, both of which complicate accurate and reproducible detection of tRNA molecules [14, 39, 40]. It is therefore not surprising that little is known about specific regulation of tRNA genes in their chromatin context. In order to overcome this knowledge gap and to circumvent the technical challenges mentioned above, we employed Epi-Decoder to delineate the local chromatin proteome of a single tRNA gene in budding yeast in a direct and unbiased manner. In Epi-Decoder, the local chromatin abundance of each protein in the cell at the barcoded locus of interest can be measured by chromatin immunoprecipitation followed by DNA-barcode sequencing and counting [41–43]. This method is sensitive and quantitative, and overcomes common hurdles associated with other proteomics approaches such as capture of a locus combined with mass spectrometry [44–46].

Here we used Epi-Decoder to delineate the proteome of a single tRNA gene, *tP(UGG)M*, in *Saccharomyces cerevisiae* to uncover chromatin-associated regulators of RNAPIII. Comparing the proteome in different metabolic conditions revealed reprogramming of the core RNAPIII transcription machinery, supporting a model of competitive binding between RNAPIII and TFIIIC. In addition, in conditions of nutrient stress we observed increased binding of known factors such as heat shock proteins, RNA processing factors, chromatin remodeling complexes but we also identified the uncharacterized factor Ykr011c/Fpt1. Deletion of *FPT1* compromised the nutrient-dependent reprogramming of RNAPIII, providing evidence for a chromatin-associated regulatory mechanism of RNAPIII assembly at tRNA genes. We expect that our dataset on tDNA proteome factors and how they change in conditions of nutrient stress will provide a valuable resource for further studies on the impact of chromatin-associated factors on tDNA transcription.

## RESULTS

### Decoding the chromatin proteome of a single native tDNA locus

To delineate the local chromatin proteome of a single tRNA gene and identify potential chromatin-associated RNAPIII regulators, we employed Epi-Decoder (as outlined in **Figure 1A**) [41–43]. Many tRNA genes in yeast are flanked by transposons or transposon LTR remnants since retrotransposons in yeast preferentially integrate near tRNA genes [47–49]. To capture a representative native chromatin environment, we employed Epi-Decoder on a tRNA gene, *tP(UGG)M*, flanked by the *YMLWTy1-2* retrotransposon. Using CRISPR/Cas9, a barcoded yeast library was generated by inserting in thousands of yeast strains a DNA-barcode between the divergent tRNA and Ty1 genes. With high-throughput synthetic genetic array (SGA) methods [50], the barcode library was crossed with a genome-wide protein TAP-tag library to create an Epi-Decoder library. In this library, each clone contained a unique barcode/protein TAP-tag combination. To account for individual barcode effects, each TAP-tagged protein was combined with three different DNA-barcodes. The three Epi-Decoder libraries were each pooled into single flasks, followed by cross-linking, chromatin shearing, chromatin immunoprecipitation (ChIP), DNA-barcode amplification and sequencing, and barcode counting. Barcode counts (ChIP versus input) served as a readout for occupancy of each tagged protein at the barcoded tDNA-Ty1 locus.

**Figure 1.**
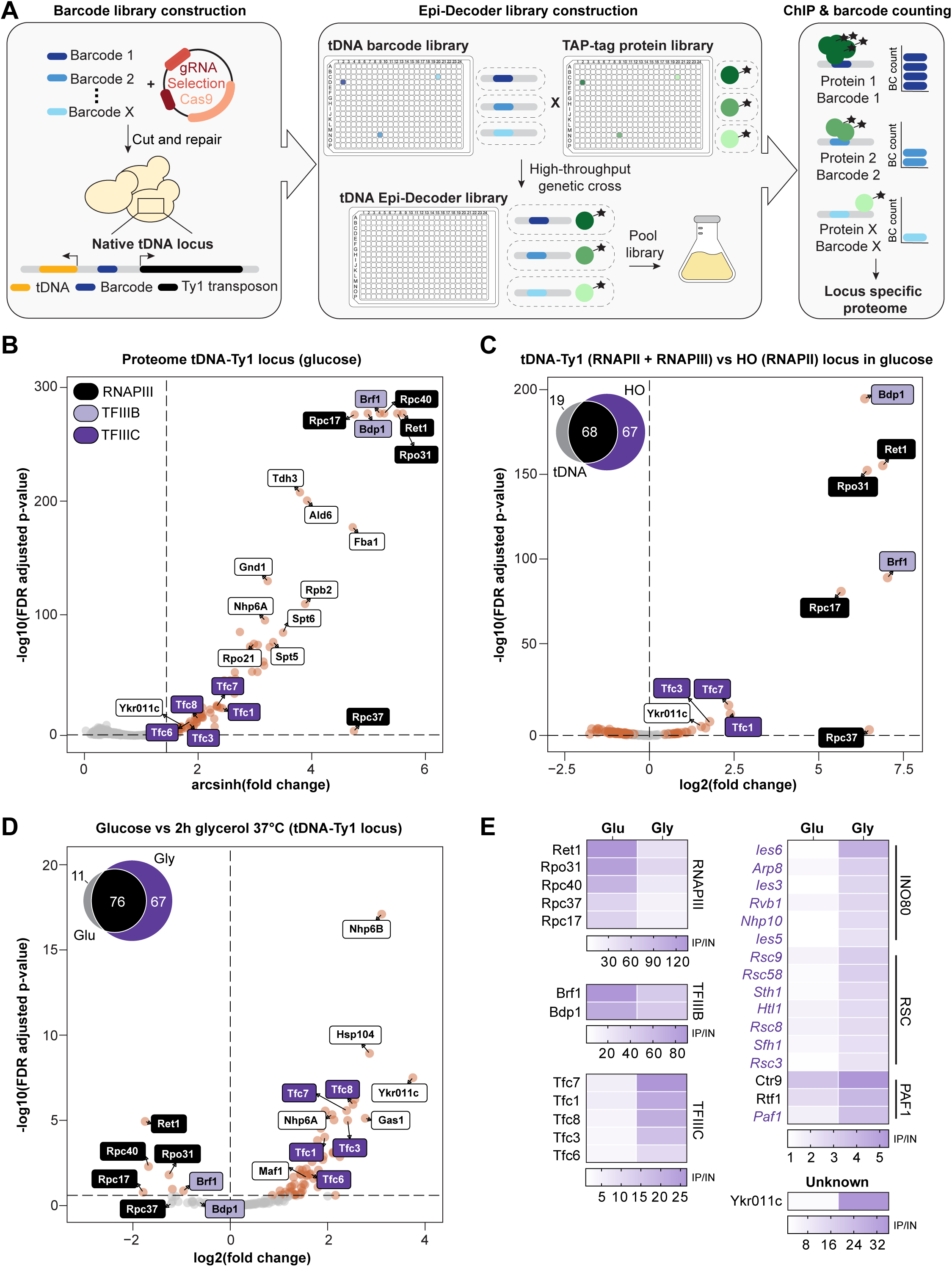
Epi-Decoder reveals rewiring of the proteome of a tDNA locus in response to nutrient availability. **(A)** Schematic overview of Epi-Decoder. In brief, a DNA-barcode repair template library and CRISPR-Cas9 construct targeting a locus of interest are transformed into yeast cells. A library of transformants containing a random 16 bp DNA-barcode at the locus of interest is arrayed, decoded, and crossed with an arrayed TAP-tag protein library in which each clone contains a different protein tagged. The resulting Epi-Decoder library contains unique combinations of barcodes and tagged proteins. After pooling the Epi-Decoder library, ChIP is performed and DNA-barcodes from ChIP and input are amplified, sequenced, and counted to provide a binding score (ChIP/input) at the locus for all proteins in the library. **(B)** Epi-Decoder scores for 3707 proteins at the tDNA-Ty1 locus in glucose; arcsinh (fold change ChIP vs input) and FDR (p-value). Subunits that belong to RNAPIII, TFIIIB or TFIIIC are color-coded. Data describes three biological replicates (different DNA-barcode protein-TAG combinations) and three technical replicates (same DNA-barcode). Proteins classified as ‘binder’ are indicated with colored dots and significance thresholds with dashed lines (basemean ≥ 400, FDR ≤ 0.01 and log_2_ fold change ≥ 1). **(C)** Binding scores of tDNA-Ty1 compared to HO locus: log_2_ (fold change ChIP_HO_ vs ChIP_tDNA-Ty1_) and FDR (p-value) for 145 factors that are ‘binder’ at either of the loci. The Euler diagram shows the number of proteins classified as ‘binder’ at the tDNA-Ty1 or HO locus. Data describes three biological replicates as described in (B). Colored dots show significant proteins and dashed lines show significance thresholds (FDR ≤ 0.25). **(D)** As in (C), fold change (log_2_ ChIP_glucose_ vs ChIP_glycerol_) between glucose and glycerol 2h (37°C) and FDR (p-value) for 154 ‘binders’ at the tDNA-Ty1 locus in either growth condition. The Euler diagram shows the number of proteins classified as ‘binder’ at the tDNA-Ty1 locus in the two conditions. **(E)** Heat maps show fold change values (ChIP vs input) for different protein complexes or families at the tDNA-Ty1 locus. The average ChIP/input of three biological replicates is shown. Protein names are color-coded based on the Euler diagram in (D). Black: shared ‘binders’ between glucose and glycerol 2h (37°C). Purple and italics: ‘binders’ in glycerol 2h (37°C). Note that the scale of each heat map is different due to large differences in cross-linking efficiency between different protein categories.

Decoding the tDNA-Ty1 proteome revealed a wide variety of proteins cross-linking to the locus. Proteins classified as ‘binder’ (base mean ≥ 400, FDR ≤ 0.01 & log_2_ fold change ≥ 1) included all known RNAPIII subunits and transcription factors present in our library, RNAPII transcription and elongation factors, and metabolic factors (**Figure 1B, Supplemental Table S1 and S2**). Histones were also observed but excluded from the analyses for technical reasons (see Supplemental Material & Methods). Because the barcoded locus contained a tDNA transcribed by RNAPIII and Ty1 retrotransposon transcribed by RNAPII and because tDNAs are hotspots for non-specific cross-linking of proteins [38], we first wanted to determine the specificity of the binders. To this end, using the same workflow, we compared binders at the tDNA-Ty1 locus (n = 87) with binders at the HO locus (n = 135), a previously decoded RNAPII promoter region [42]. The majority of proteins (n = 68) occupied both the tDNA-Ty1 and HO locus, including RNAPII transcription and elongation factors, metabolic factors, chromatin remodelers (e.g. FACT, INO80, RSC and PAF1 complexes) and RNA processing factors (**Figure 1C, S1A, S1C, Supplemental Table S1, S3 and S4**). One major advantage of Epi-Decoder is that binding of each protein is interrogated in a single pool with all other proteins. As a result, potential non-specific cross-linking is expected to be eliminated by internal normalization. Indeed, specific tDNA binders included subunits of the RNAPIII, TFIIIB and TFIIIC complexes suggesting that the tDNA-Ty1 binder set is not affected by potential hotspot artifacts (**Figure 1C**).

### The tDNA proteome is dynamically regulated in response to nutrient availability

In nutrient-rich conditions (**Figure 1B**), tRNA genes are actively transcribed by RNAPIII, but in repressive conditions, the repressor Maf1 is localized to the nucleus and tDNA transcription is restrained. Maf1 prevents the assembly of RNAPIII and TFIIIB onto tDNAs but is itself not a component of the tDNA chromatin and therefore considered a trans-factor [51]. To investigate whether specific chromatin-associated regulators of tDNAs exist, we applied Epi-Decoder in a condition of repressed tDNA transcription. After cells were grown to mid-log phase in media containing glucose at 30°C, cells were switched to glycerol at 37°C for 2 hours, a condition associated with increased Maf1 activity and repressed RNAPIII transcription [5, 24]. Differential analysis showed extensive reprogramming of the general RNAPIII transcription factors upon a switch to repressive conditions (**Figure 1D, 1E, S1D, Supplemental Table S1, S4 and S5**). Binding of both RNAPIII and TFIIIB was decreased, corroborating with repressed tDNA transcription. In contrast, binding of TFIIIC was increased. Previous studies on individual subunits of these complexes have suggested that RNAPIII and TFIIIC are inversely correlated due to their competitive binding for the same locus [52, 53]. Here we confirm and extend this proposed model from the level of individual subunits to their protein complexes. Unexpectedly, the repressed state of the tDNA-Ty1 locus showed an overall increased number of binders (n = 143) compared to the active state (**Figure S1B**). In repressive conditions, relative binding of the two high-mobility group proteins Nhp6a and Nhp6b, known to aid tDNA transcription [54–56], was increased. In addition, we observed an increased occupancy of the chromatin remodeling complexes INO80, RSC and the RNAPII elongation complex PAF1, suggesting a broad chromatin adaptation to repressive conditions (**1E, S1D,** note that the scale of each heat map is unique due to large differences in cross-linking efficiency between different protein complexes). Since the occupancy of RNAPII subunits was not consistently altered in repressive conditions, the observed changes were most likely not caused by altered transcription of the proximal Ty1 element (**Figure S1D**). Additionally, we observed increased relative occupancy of heat shock proteins, important chaperones that maintain protein homeostasis, and RNA processing factors. Lastly, we observed increased binding of several transcriptional repressors, suggesting an overall repressed chromatin-state (**Figure S1D**). Overall, the dynamic proteome atlas of the tDNA-Ty1 locus provides a valuable resource for identifying putative chromatin-associated factors that potentially regulate tDNA biology.

### A novel factor enriched in the proteome of tRNA genes

Having determined that the tDNA chromatin proteome is reprogrammed in changing metabolic conditions, we next asked whether candidate factors are embedded in the proteome that might have a regulatory function. Focusing on binders that show dynamic chromatin interactions, the protein Ykr011c drew our attention. Ykr011c was found at the tDNA-Ty1 locus but not the reference HO promoter locus (**Figure 1C**) and showed substantially increased binding in repressive conditions (**Figure 1D**). This specific binding and dynamic behavior suggested that Ykr011c could be a bona fide but previously unknown member of the tDNA chromatin proteome. Ykr011c is a protein of unknown function and structure, and phylogenetic analysis showed that this protein is conserved within the Saccharomycetales clade (**Figure S1E**). To validate the Epi-Decoder results, we first determined the genome-wide binding profile of Ykr011c by ChIP-sequencing. An RNAPIII subunit (Rpo31) and RNAPII subunit (Rpb2) were taken along as controls for RNAPIII and RNAPII transcribed regions, respectively. We also included a TAP-tagged ribosomal protein (Rpl13a) as a negative control because tRNA genes have been reported to act as sites of non-specific binding [38]. Visual inspection of the ChIP-seq data at the tDNA-Ty1 Epi-Decoder locus showed that Ykr011c is enriched at the *tP(UGG)M* tRNA gene and absent from the Ty1 element (**Figure 2A**). At a genome-wide level, Ykr011c was found to uniquely bind all nuclear tRNA genes and a few other RNAPIII-regulated genes, including *RPR1, SNR6,* and *SCR1* and the tDNA relics *ZOD1* and *iYGR033c* (**Figure 2B, S2A**). We found no evidence for Ykr011c binding at ETC loci, sites in the genome previously described as regions of extra TFIIIC (ETC) binding [57] (**Figure S2A**). These results confirmed binding of Ykr011c to the tDNA-Ty1 locus and furthermore showed that Ykr011c is a novel specific tDNA-binding protein. From this point onward we will refer to Ykr011c as Fpt1 (Factor in the Proteome of tDNAs number 1).

**Figure 2.**
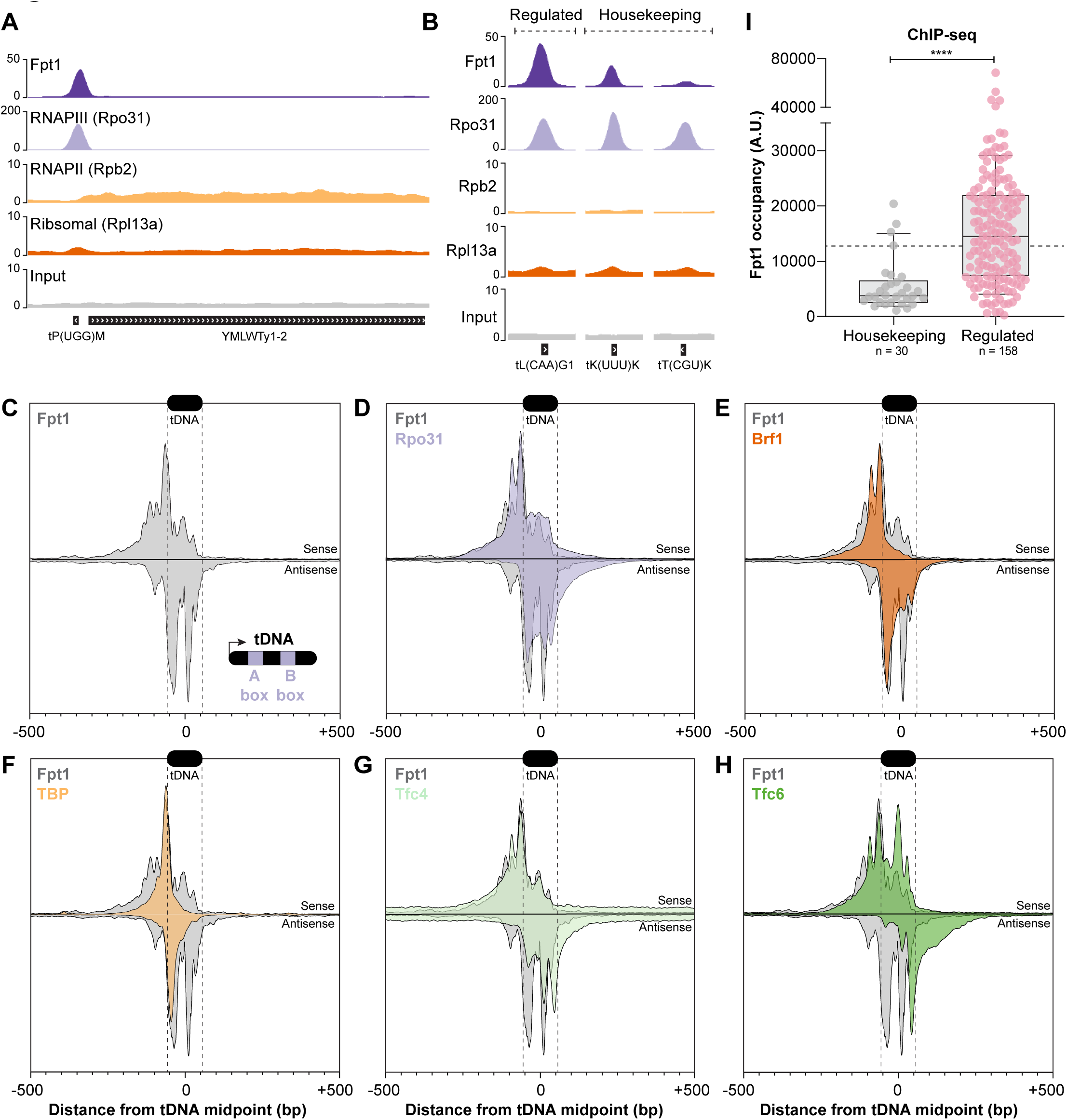
Fpt1 binds uniquely to RNAPIII transcribed genes and is enriched at regulated tDNAs. **(A)** Genome tracks show pooled ChIP-seq data of three biological replicates for TAP-tagged Fpt1, Rpo31, Rpb2 and Rpl13a at the Epi-Decoder locus: *tP(UGG)M* tRNA gene and *YMLWTy1-2* retrotransposon. **(B)** As in (A), tRNA genes *tL(CAA)G1, tK(UUU)K* and *tT(CGU)K.* **(C)** ChIP-exo metagene profile of Fpt1 at 261 tRNA genes from a single representative ChIP-exo experiment. Data is shown for a 1 kb window with the tDNA midpoint at zero. Plots of ChIP-exo tag 5’ ends (exonuclease stop sites) are inverted for tags mapping to the non-transcribed (anti-sense) strand. The y-axis shows linear arbitrary units (AU) which are not equal across plotted datasets. The black box at the top shows the average position of a representative tRNA gene, start and end are indicated with dashed lines. The insert in the lower right corner shows a schematic overview of a tRNA gene, containing the internal A- and B-box promoter elements. **(D-H)** As in (C), ChIP-exo metagene profiles of Rpo31, Brf1, TBP, Tfc4 and Tfc6 overlaid with Fpt1. ChIP-exo data on the RNAPIII subunits and transcription factors is described in [58]. **(I)** Fpt1 occupancy (ChIP-seq) at housekeeping (n = 30) and regulated (n = 158) tRNA genes. The used classification of tRNA genes is described in [25] and in the Supplemental Material & Methods. The average Fpt1 occupancy (13142) across 243 tRNA genes (see Supplemental Material & Methods) is indicated with a dotted horizontal line. Significance was determined using a Welch-corrected unpaired two-tailed t-test, ****: p < 0.0001.

To analyze the binding of Fpt1 to tDNAs at higher resolution, we applied ChIP-exo to provide detailed information on the protein-DNA contacts that Fpt1 makes [58, 59]. ChIP-exo confirmed the binding of Fpt1 to tRNA genes observed by ChIP-seq (**Figure 2C**). In addition, the tDNA metagene ChIP-exo profile of Fpt1 showed several contact points, both upstream and within the tRNA gene body. Superimposing the ChIP-exo profile of Fpt1 with those of known RNAPIII factors revealed shared tDNA contact points, especially with TFIIIB and TFIIIC (**Figure 2D-H**). This suggests that Fpt1 makes contact with tDNAs at least in part through the core RNAPIII transcription machinery complex.

Unexpectedly, in contrast to RNAPIII, which was distributed relatively evenly across all tRNA genes, ChIP-seq and ChIP-exo analysis displayed varied levels of Fpt1 binding (**Figure 2B, S2A-C, Supplemental Table S6**). While some tRNA genes were bound by high levels of Fpt1, others were scarcely occupied. To investigate whether this could reflect a functional difference, we compared Fpt1 binding at tRNA genes with the previously determined distribution of transcriptionally engaged RNAPIII at tRNA genes in response to changing nutrient conditions [25]. Turowksi et al. used UV cross-linking and analysis of cDNA (CRAC) to capture nascent RNAs bound by RNAPIII and identified two groups of tRNA genes (**Supplemental Table S6**, and see Supplemental Material & Methods). One canonical group of ‘regulated tRNA genes’ showed efficient eviction of engaged RNAPIII upon a switch from glucose to glycerol (low RNAPIII retention score) and this repression was dependent on Maf1 (high Maf1 dependency score). Another group of ‘housekeeping tRNA genes’ was less responsive to changing nutrients (high RNAPIII retention score) and less affected by loss of Maf1 (low Maf1 dependency score). Inspection of Fpt1 binding at tRNA genes ranked by their CRAC response score suggested that Fpt1 binding is on average higher at regulated tRNA genes and lower (but still higher than background, **Figure 2B, S2A, Supplemental Table S6**) at housekeeping tRNA genes (**Figure 2I**). Indeed, Fpt1 occupancy negatively correlated with retention of engaged RNAPIII upon a switch from glucose to glycerol and positively correlated with Maf1-dependent repression (**Figure S2D-G**). The correlation between Fpt1 occupancy and efficiency of RNAPIII regulation suggests that Fpt1 may be part of a chromatin-associated mechanism of tRNA gene regulation.

### Fpt1 responds to changing nutrient availability

To investigate the role of Fpt1 in tDNA regulation, we first examined Fpt1 binding at several tDNA loci, representing the classes housekeeping (*tP(UGG)M*, *tR(CCG)L*, *tK(UUU)K*, *tL(CAA)G1*) and regulated (*tL(CAA)G1*, tP*(UGG)A*, *tM(CAU)E*, *tV(CAC)D*) in conditions of repressed RNAPIII activity. Cells were grown to mid-log phase in glucose and subsequently subjected to two independent repressive conditions by switching to ethanol (30°C) or glycerol (37°C) as a carbon source. In both conditions, RNAPIII activity is repressed, but glycerol 37°C has frequently been used since Maf1 is essential in this condition (**Figure S6C** and [24]). Fpt1 occupancy (defined as ChIP over input) increased at all tested tRNA genes in repressive conditions and is on average (n = 8 tRNA genes) largest in the 2h glycerol condition (14.8-fold increase), followed by 2h ethanol (8.3-fold increase) and 15 min glycerol (5.0-fold increase) (**Figure 3A**). These results corroborate the nutrient response observed in Epi-Decoder and demonstrate that increased Fpt1 occupancy occurs at other tRNA genes as well. Additionally, a similar increase in Fpt1 binding was observed for the tDNA relic *ZOD1* and RNAPIII regulated non-tRNA genes *SNR6, SCR1* and *RPR1* (**Figure S3A**).

**Figure 3.**
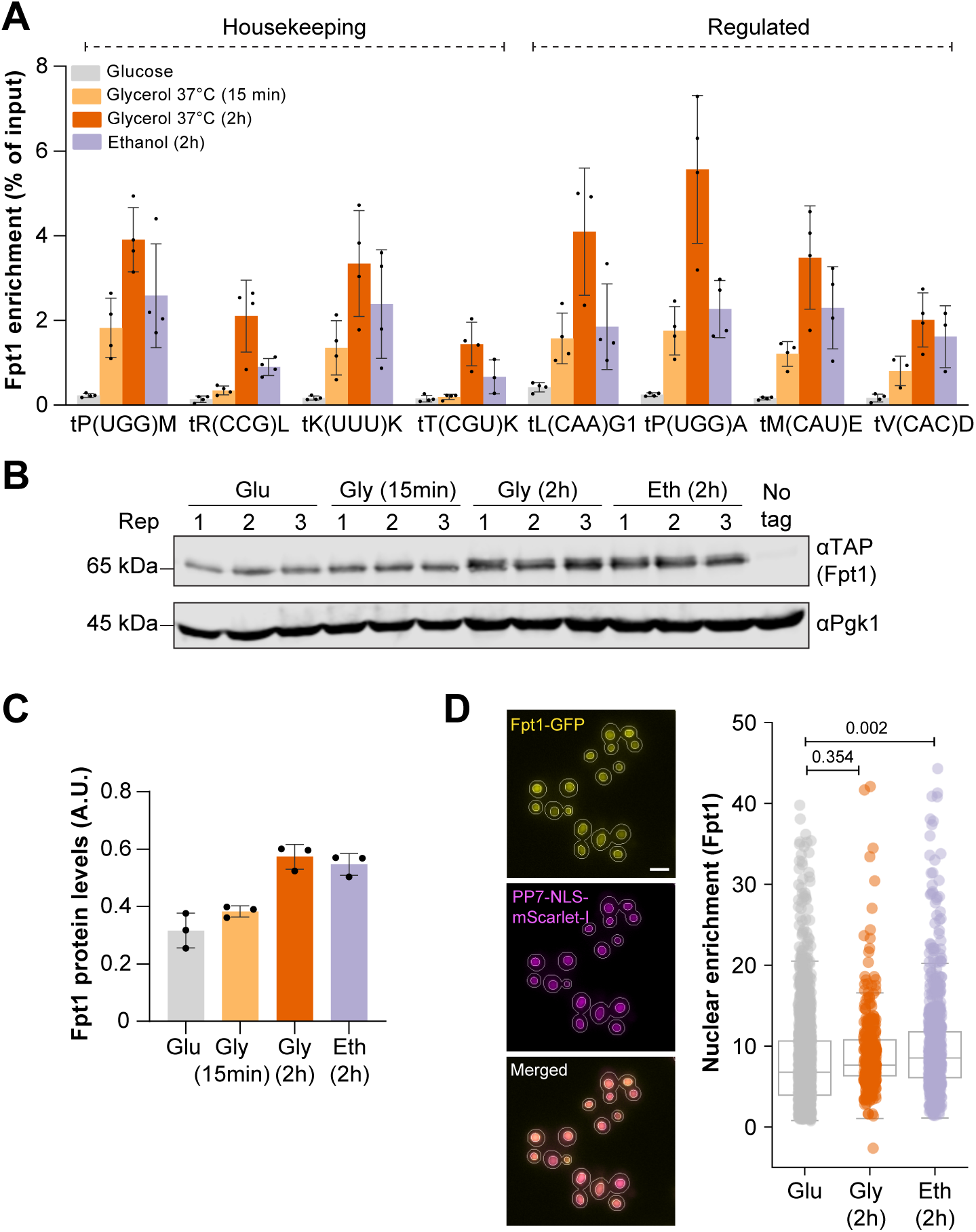
Fpt1 responds to changing nutrient availability. **(A)** Fpt1 enrichment (ChIP) at different tRNA genes (housekeeping and regulated subsets are indicated) in glucose, glycerol 2h and 15 min (37°C), and ethanol 2h. The average and standard deviation of four biological replicates is shown (unless a replicate was excluded from the qPCR data due to technical variation). **(B)** Fpt1-TAP immunoblot (anti-TAP) of three biological replicates in glucose, glycerol 2h and 15 min (37°C), and ethanol 2h. Pgk1 was used as a loading control. No tag is a negative control strain lacking a TAP-tag (BY4741). **(C)** Quantification of (B). **(D)** Left, representative images of Fpt1 localization in live yeast cells growing in glucose. From top to bottom: Fpt1-GFP (yellow), PP7-NLS-mScarlet-I as a nuclear marker (magenta) and the merged channel. Cellular masks to locate cells and nuclei are indicated with white lines. Scale bar: 3 µm. Right, quantification of Fpt1 nuclear enrichment, defined as the nuclear median intensity divided by the total median intensity (see Figure S3B-D), of three biological replicates in glucose (n = 1536 cells), glycerol 2h 37°C (n = 327 cells), and ethanol 2h (n = 771 cells). Circles show data for individual cells and box plots show the distribution of the data where the box indicates the quartiles and whiskers extend to show the distribution, except for outliers. The p-values depicted are determined based on bootstrapping [73].

To determine whether increased Fpt1 binding to tDNAs during repressive conditions was caused by increased Fpt1 protein expression, we measured global cellular protein levels of Fpt1 in response to changing nutrient conditions. Immunoblotting showed that increased Fpt1 occupancy at tRNA genes was accompanied by an increase in Fpt1 protein levels upon a switch to repressive conditions (**Figure 3B-C**). However, the increase in Fpt1 occupancy at tRNA genes (14.8-fold in 2h glycerol 37°C) is not equivalent to the increase in protein level (1.7-fold in 2h glycerol 37°C) suggesting that additional mechanisms play a role in recruiting Fpt1 to chromatin. We also determined the cellular localization of Fpt1 in nutrient-rich and repressive conditions. GFP-tagged Fpt1 localized to the nucleus in all tested conditions and showed only a modest increase in nuclear enrichment in repressive conditions (**Figure 3D, S3B-D**). This suggests that, in contrast to Maf1 [60], Fpt1 is constitutively present in the nucleus and activated by different mechanisms. Altogether, the increased tDNA occupancy and protein levels of Fpt1 in conditions of repressed RNAPIII transcription are in agreement with a potential regulatory role at tDNAs.

### Deletion of Fpt1 compromises eviction of RNAPIII upon stress

Studying tDNA transcription poses multiple challenges due to the stable structure and many base modifications of tRNA molecules that interfere with reverse transcription and detection by sequencing. In addition, while identical tRNA genes across the genome can have different properties, the identical mature tRNA sequences derived from these different genomic locations cannot be uniquely mapped to each of the originating tDNA loci. To circumvent these issues, we focused on RNAPIII occupancy at the chromatin and how it is affected by Fpt1. To this end, we performed ChIP-qPCR of the largest RNAPIII subunit (Rpo31) in both wild type and *fpt1*Δ strains. Cells were grown to mid-log phase in glucose and shifted to repressive conditions for 2 hours. A stress response of RNAPIII in wild-type cells could be observed at all tRNA genes examined and for most tRNA genes RNAPIII loss could already be observed after 15 minutes in glycerol (37°C) (**Figure 4A**). On the contrary, RNAPIII occupancy at *ZOD1, SNR6, SCR1* and *RPR1* was less affected by repressive conditions suggesting a different mechanism of regulation [24] (**Figure S4A**). Decreased Rpo31 occupancy in all tested conditions in both wild type and *fpt1*Δ was on average most profound at tRNA genes belonging to the regulated tDNA subset (n = 4) (**Figure 4B, S4B**), which is in agreement with previous observations [25], although the response was variable among tRNA genes within each class. Looking at the role of Fpt1, we observed that in the absence of Fpt1, more Rpo31 was retained at tRNA genes compared to wild type (**Figure 4C-F, S4C-F**). The gain of Rpo31 occupancy at tRNA genes in *fpt1*Δ suggests that Fpt1, directly or indirectly, regulates RNAPIII eviction in all tested conditions (**Figure S4G**). The extent to which Rpo31 occupancy increased in *fpt1*Δ was variable and distinct across tRNA genes but correlated with the two tRNA gene classes suggesting that regulated tRNA genes are more dependent on Fpt1 (**Figure 4G**). To determine whether the increase in Rpo31 occupancy in *fpt1*Δ was caused by altered Rpo31 protein levels, we performed immunoblotting. In wild-type cells, Rpo31 protein levels decreased in repressive conditions (**Figure 4H-I, S4H**), which is in agreement with decreased Rpo31 occupancy at the tRNA genes (**Figure 4A**) and the previously observed proteasomal degradation in stress [61]. The reduction in Rpo31 protein levels was similar in *fpt1*Δ cells (**Figure 4I**), suggesting that the increased Rpo31 occupancy at tDNAs in *fpt1Δ* is not a consequence of increased Rpo31 protein levels.

**Figure 4.**
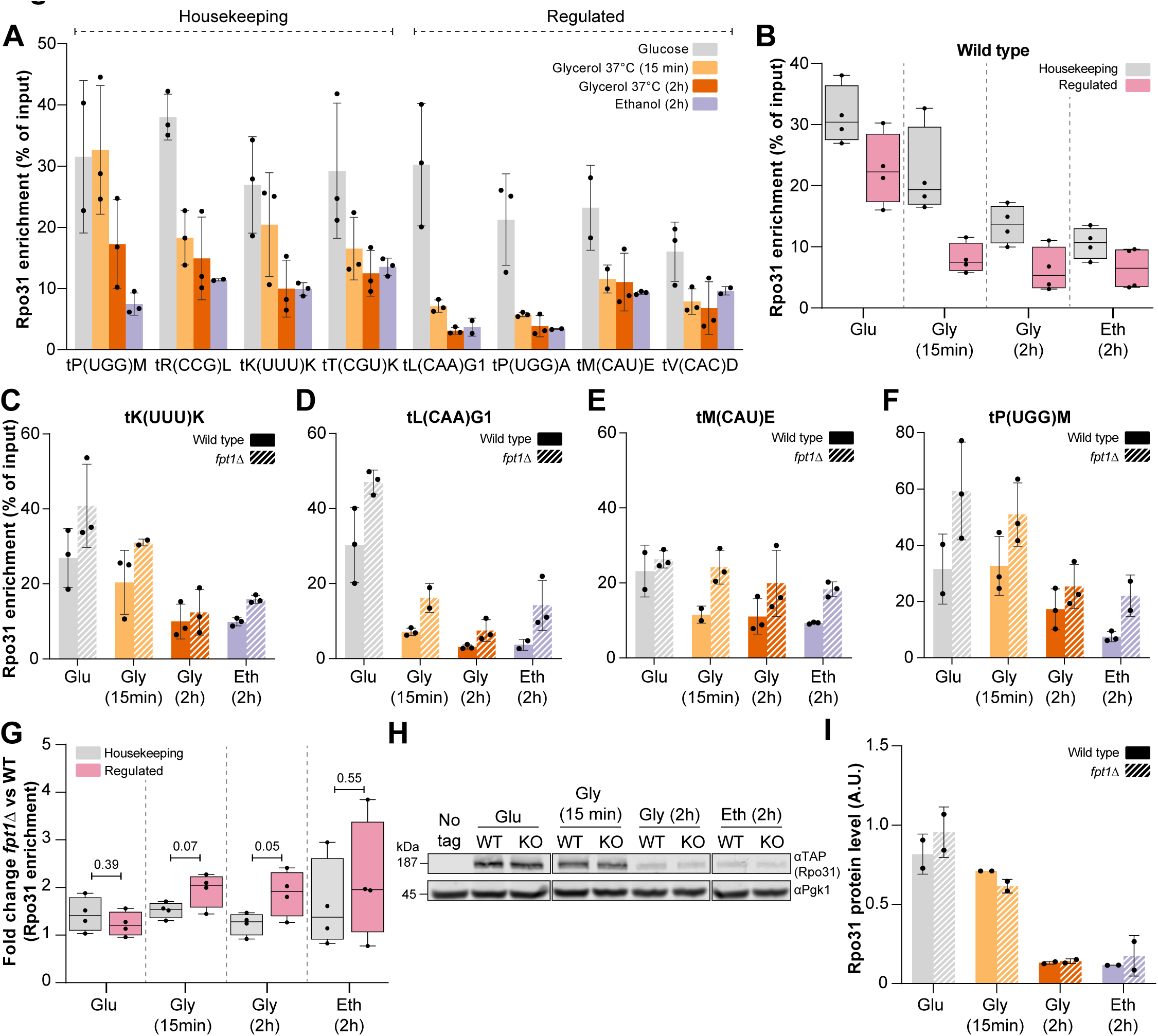
Deletion of Fpt1 compromises eviction of RNAPIII upon nutrient perturbation. **(A)** Rpo31 enrichment (% of input) at different tRNA genes (divided in housekeeping and regulated subsets) in glucose, glycerol 2h or 15 min (37°C) and ethanol 2h. The average and standard deviation of three biological replicates is shown (unless a replicate was excluded from the qPCR data due to technical variation). **(B)** Rpo31 enrichment (% of input) in wild type at housekeeping tRNA genes (grey, n = 4) and regulated tRNA genes (pink, n = 4) in glucose, glycerol 2h and 15 min (37°C), and ethanol 2h. For each tRNA gene, the average of three biological replicates is shown. Whiskers indicate the minimum and maximum. **(C-F)** Rpo31 enrichment (% of input) in wild type and *fpt1Δ* at different tRNA genes in glucose, glycerol 2h and 15 min (37°C), and ethanol 2h. The average and standard deviation of three biological replicates is shown (unless a replicate was excluded from the qPCR data due to technical variation). **(G)** Fold change between wild type and *fpt1Δ* for Rpo31 enrichment (% of input) at housekeeping tRNA genes (grey, n = 4) and regulated tRNA genes (pink, n = 4). For each tRNA gene, the average of three biological replicates is shown. Whiskers indicate the minimum and maximum. Significance was determined using an unpaired two-tailed t-test. **(H)** Rpo31-TAP (anti-TAP) immunoblot in glucose, glycerol 2h and 15 min (37°C), and ethanol 2h. WT: wild type, KO: *fpt1Δ*. Pgk1 was used as a loading control. No tag is a negative control strain lacking a TAP-tag (BY4741). **(I)** Quantification of (H) and S4H showing the average and standard deviation of two biological replicates.

### Fpt1 functions as a central regulator of the RNAPIII transcription machinery

The partial eviction of Rpo31 in *fpt1Δ* raises the question to what extent the full RNAPIII complex and other members of the general tDNA transcription machinery are affected. To address this question, we took advantage of Epi-Decoder to study binding changes at the barcoded tDNA locus of all proteins in parallel in wild type compared to *fpt1Δ*. We repeated Epi-Decoder after crossing into the library an *fpt1Δ* allele as outlined in **Figure 5A**. Differential analysis was done on binders (base mean ≥ 400, FDR ≤ 0.01 & log_2_ fold change ≥ 1) in glucose (n = 87 wild type, n = 77 *fpt1*Δ, **Figure S5A, Supplemental Table S1, S7**) and 2h glycerol 37°C (n = 143 wild type, n = 138 *fpt1*Δ, **Figure S5B, Supplemental Table S1, S5**). Similar to wild type, the majority of proteome-members included chromatin remodelers, transcription factors, metabolic enzymes, heat shock proteins and RNA processing factors (**Figure 5B**). In conditions of active transcription, *FPT1* deletion showed mild effects on the relative occupancy of RNAPIII, TFIIIB and TFIIIC. This effect was enhanced when cells were subjected to repressive conditions (**Figure 5C**). In *fpt1Δ,* we observed a partial eviction of RNAPIII and TFIIIB subunits compared to wild type. On the contrary, binding of TFIIIC subunits was less increased in *fpt1Δ.* Additionally, in repressive conditions *fpt1Δ* caused decreased relative binding of several heat shock factors (**Figure 5C**). Other factors such as RNAPII subunits (regulating the proximal Ty1 element) and chromatin remodelers remained largely unaffected (**Figure 5C, S5C**). The observed coupling between increased RNAPIII and TFIIIB and decreased TFIIIC occupancy provides evidence for a competitive-binding model that has previously been proposed [52, 53].

**Figure 5.**
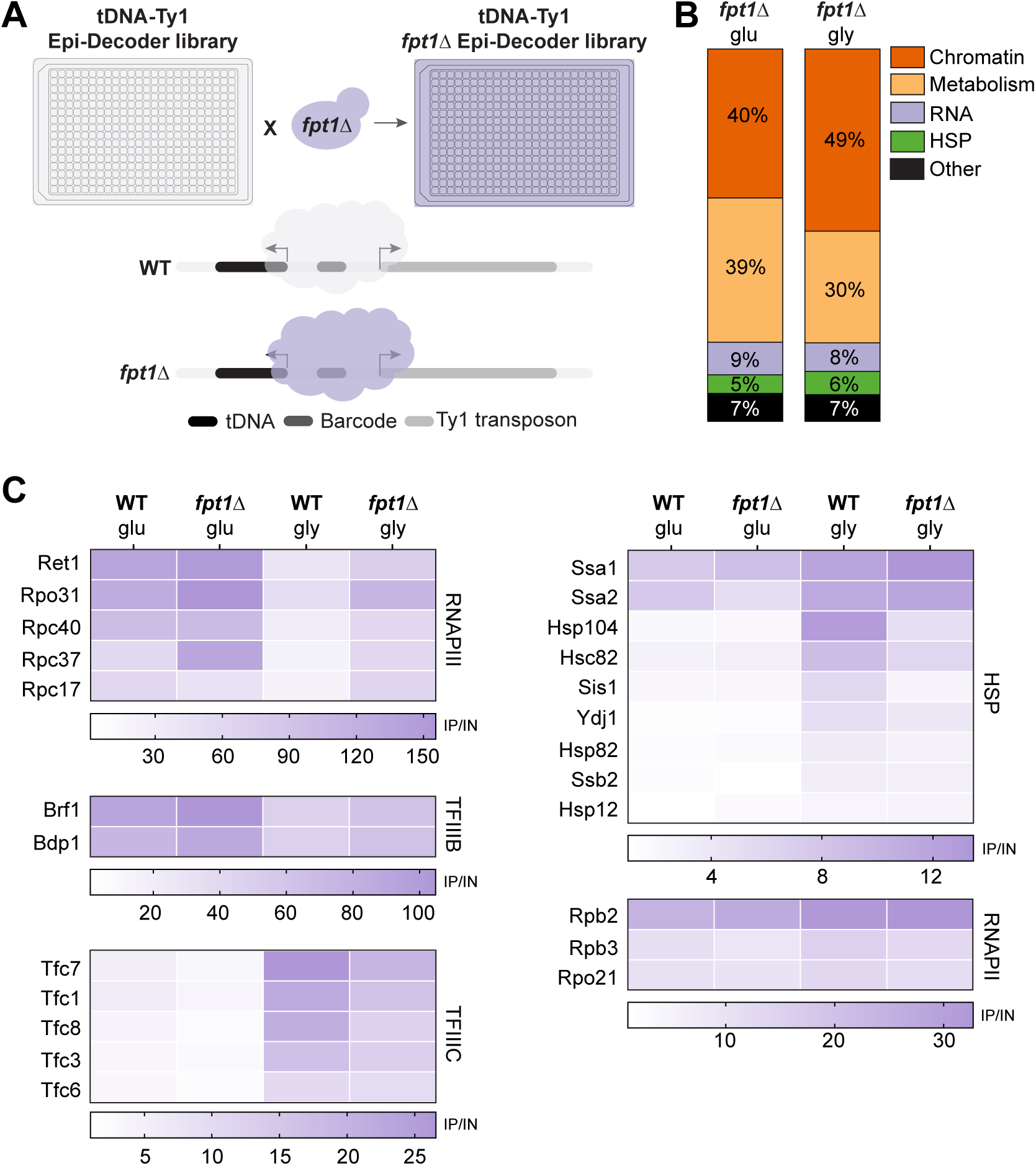
Fpt1, embedded in the tDNA proteome, functions as a regulator of the RNAPIII transcription machinery. **(A)** Schematic outline: an *FPT1* knockout strain is crossed with the wild-type Epi-Decoder library using SGA, resulting in an *fpt1Δ* Epi-Decoder library and a tDNA proteome that can be compared to the original wild-type proteome. **(B)** Fraction of proteins classified as ‘binder’ in *fpt1Δ* glucose or glycerol 2h (37°C) for different functional classes: chromatin, metabolism, RNA, heat shock proteins (HSP) and others. **(C)** Heat maps show fold change values (ChIP vs input) at the tDNA-Ty1 locus for different protein complexes or families for wild type or *fpt1Δ* in glucose and 2h glycerol (37°C). WT = wild type, KO = *fpt1Δ*, glu = glucose, gly = glycerol 2h (37°C). The average ChIP/input of three biological replicates is shown. Note that the scale of each heat map is different due to large differences in cross-linking efficiency between different protein categories.

### Fpt1 affects tuning of ribosome biogenesis genes and cellular fitness in repressive conditions

Our data suggest that Fpt1 is part of a previously unknown chromatin-associated mechanism, being in proximity to and regulating the dynamic assembly of the RNAPIII transcription machinery at tRNA genes. To explore how the defect in RNAPIII dynamics caused by *fpt1Δ* is linked to general cell physiology, we performed mRNA-sequencing in wild type and *fpt1*Δ. Cells were grown to mid-log phase in glucose containing media and subsequently switched to a non-fermentable carbon source (ethanol) for 2 hours at 30°C to avoid excessive temperature effects on the transcriptome. After 2 hours in ethanol, a large fraction of genes was differentially expressed in wild-type cells (**Figure 6A, Supplemental Table S8**). As expected, genes involved in ribosome biogenesis (RiBi), gene expression, and metabolic processes were downregulated, while genes involved in mitochondrial translation, aerobic respiration and oxidation were upregulated (**Figure S6A-B, Supplemental Table S8**). We did not observe many differentially expressed genes between wild type and *fpt1*Δ in glucose (**Figure 6B, Supplemental Table S8**) but upon a switch to repressive conditions over 400 genes were differentially expressed (**Figure 6C, Supplemental Table S8**). Gene ontology analysis of the 313 upregulated genes relative to wild type showed enrichment of 97 RiBi genes (as defined in [62]) in *fpt1*Δ (**Figure 6D, Supplemental Table S8**), suggesting that shutdown of RiBi gene expression was dysregulated in *fpt1*Δ. Fpt1 uniquely binds RNAPIII regulated genes (and not RNAPII target genes) and its absence leads to reprogramming of the tDNA proteome in repressive conditions. Therefore, our results suggest that Fpt1 indirectly affects transcription of ribosome biogenesis genes most likely through its role in RNAPIII assembly at tDNAs. This model is supported by the observation that strains expressing a mutant form of the RNAPIII catalytic subunit Rpc128 also show a weak increase in RiBi gene expression [63].

**Figure 6.**
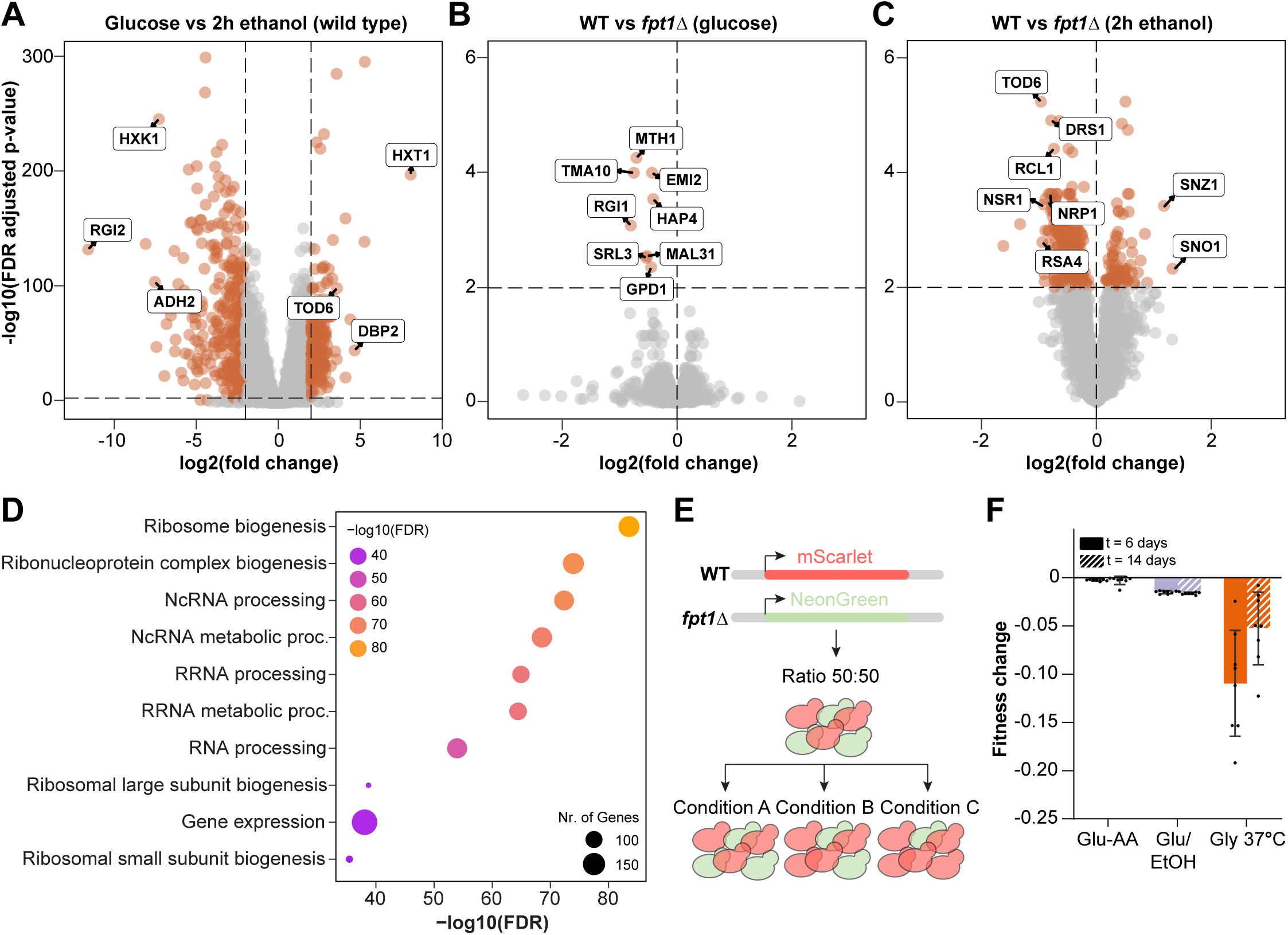
Fpt1 affects tuning of ribosome biogenesis genes and cellular fitness in repressive conditions. **(A)** Differential mRNA expression in glucose and ethanol (2h) in wild type: log_2_ fold change and FDR (p-value) for 5074 expressed genes. Colored dots represent significant differentially expressed genes (FDR ≤ 0.01 & log_2_ fold change ≥ 2). Upregulated genes in ethanol 2h (n = 1929) are depicted on the left, upregulated genes in glucose (n = 2026) are depicted on the right. RNA-sequencing data represents three biological replicates for wild type and four biological replicates for *fpt1Δ*. **(B)** Differential gene expression in wild type and *fpt1Δ* in glucose: log_2_ fold change and FDR (p-value) for 5080 expressed genes. Colored dots represent significant differentially expressed genes (FDR ≤ 0.01). Upregulated genes in *fpt1Δ* compared to wild type are depicted on the left (n = 8). **(C)** As in (B), differential gene expression in wild type and *fpt1Δ* in ethanol 2h: log_2_ fold change and FDR (p-value) for 4978 expressed genes. Upregulated genes in *fpt1Δ* compared to wild type are depicted on the left (n = 313) and downregulated genes on the right (n = 92). **(D)** Gene ontology analysis of genes upregulated in *fpt1Δ* in ethanol 2h (n = 313). **(E)** Schematic overview of the competitive growth assay. The fluorescent markers mScarlet and NeonGreen were inserted at a neutral intergenic locus in wild type and *fpt1Δ* in both combinations. Red and green cells were mixed in a 50:50 ratio and subjected to different growth conditions to study the effect on the ratio wild-type and *fpt1*Δ cells. **(F)** Relative fitness defect, expressed with the Malthusian coefficient, of *fpt1Δ* compared to wild type in glucose but with alternating levels of non-auxotrophic amino acids (Glu-AA), alternating carbon source (Glu/EtOH), and glycerol 37°C (Gly 37°C) at day 6 (solid fill) and day 14 (dashed fill). Data of four biological replicates including a color-swap is shown (n = 8, and see Figure S6D).

To determine the functional consequences of the observed altered RNAPIII dynamics and ribosome biogenesis gene expression, we determined the cellular fitness of *fpt1*Δ cells. Deletion of *FPT1* did not lead to observable growth defects in spot test analysis in conditions of active or repressed transcription (**Figure S6C**). As a more sensitive approach to study cellular fitness of *fpt1*Δ, we performed a competitive growth assay as outlined in **Figure 6E**, using different growth regimens (see Supplemental Material & Methods). Briefly, a different fluorescent reporter (NeonGreen or mScarlet) was inserted at a safe-harbor intergenic locus in wild-type and *fpt1*Δ cells. A color-swap was done to account for reporter effects (**Figure S6D**). At t = 0, both wild-type and *fpt1*Δ cells were equally mixed and maintained in batch culture using different growth conditions for two weeks. To include previously used repressive conditions, cells were maintained in glycerol at 37°C (Gly 37°C) or in conditions of alternating carbon sources by switching back and forth to ethanol or glucose at 30°C (Glu/EtOH). As a reference, cells were grown in glucose media but with conditions of alternating levels of non-auxotrophic amino acids (Glu-AA). Every few days, cultures were diluted to maintain a low density. Samples were taken at t = 0, t = 6 and t = 14 days to analyze the ratio of green to red cells by flow cytometry. While loss of Fpt1 did not affect fitness in glucose (Glu-AA), in conditions of alternating carbon source (Glu/EtOH) and in glycerol 37°C, *fpt1*Δ cells showed reduced competitive growth relative to wild-type cells (**Figure 6F**). These findings suggest that regulation of tDNA proteome dynamics by Fpt1 is required for optimal growth when yeast cells transiently or constitutively encounter metabolic conditions that require tuning of tDNA repression.

## DISCUSSION

Transfer-RNA genes synthesize tRNAs at high rates, but their expression is tightly tuned in response to nutrient availability. Being self-contained elements, tRNA genes have long been considered as a homogeneous group of genes regulated by general signaling pathways and mechanisms *in trans* such as Maf1. However, an alternative view has recently emerged. Indeed, the relative abundance of individual tRNAs varies considerably across tissues and cell lines, in cells engaged in proliferation or differentiation, and in pathologies such as cancer. Data from yeast and humans suggest the existence of tDNA regulatory mechanism acting at the level of chromatin but these mechanisms largely remained elusive, in part due to technical challenges associated with analyzing tDNAs and their output. Here, by taking advantage of DNA-barcode sequencing and yeast genetics (Epi-Decoder), we observe that the tDNA proteome in yeast is highly dynamic. This highly dynamic proteome contains a previously unknown chromatin-associated factor regulating the assembly of the RNAPIII transcription machinery at tRNA genes.

By comparing a tDNA proteome in active and repressive conditions, we observed a major reprogramming of the core transcriptional machinery. Previous studies showing an inverse correlation between selected subunits of TFIIIC and RNAPIII led to the model that TFIIIC and RNAPIII directly compete for binding to tDNAs [52, 53]. Here we extend and thereby confirm this model by demonstrating the same behavior for all TFIIIC and RNAPIII subunits present in our analyses, and by including TFIIIB. Further support comes from the *fpt1Δ* background, in which reprogramming of the tDNA transcription machinery was compromised, affecting RNAPIII/TFIIIB and TFIIIC in opposite ways. Importantly, our results suggest that this competitive reprogramming extends to other factors, including non-canonical tDNA binding proteins (**Figure 1E, S1D, 5C and S5C**). However, to determine whether the observed differences of non-canonical tDNA factors in Epi-Decoder are mediated by the tRNA or Ty1 gene and to understand the functional consequences of the observed proteome dynamics, further studies are required. Overall, the dynamic proteome atlas of a barcoded tDNA locus provides a valuable resource for identifying chromatin-associated factors that potentially regulate tDNA biology.

Among the proteins not previously associated with tDNAs, Epi-Decoder revealed the presence of Fpt1, a protein of unknown function. Our studies suggest that Fpt1 is a genuine member of the tDNA proteome with regulatory functions. Fpt1 binds specifically at RNAPIII-transcribed genes, its abundance is variable among tDNAs across the genome and correlates with the sensitivity to RNAPIII loss at tDNAs, and its abundance increased upon nutrient perturbation. Moreover, reprogramming of the tDNA proteome in conditions of repressed transcription is compromised in *fpt1Δ* cells, resulting in retention of RNAPIII and TFIIIB and lower levels of TFIIIC. Together, these findings point to the existence of a chromatin-associated regulatory mechanism of RNAPIII dynamics in which Fpt1 plays a role. Fpt1 seems to directly or indirectly promote eviction of RNAPIII, by mechanisms that are currently unknown. However, the mechanism seems distinct from that of Maf1. While Maf1 is actively transported to the nucleus in conditions of repressed RNAPIII transcription, Fpt1 is persistently present in the nucleus and already bound to tDNAs in conditions of active transcription. Furthermore, while Maf1 does not ChIP well to chromatin in both yeast (**Figure 1D** and [64]) and humans [31], Fpt1 is easily detectable and its abundance increases in conditions of stress. Fpt1 seems to contact tDNAs through interactions with the core RNAPIII transcription machinery. The ChIP-exo pattern of Fpt1 matched individual parts of the TFIIIB (TBP and Brf1) and TFIIIC (Tfc4 and Tfc6) patterns (**Figure S2C-H**). This correlated pattern indicates that Fpt1 is in very close proximity to both TFIIIB and TFIIIC. While TFIIIC also showed differential binding among tDNAs, it should be noted that ETCs, sites of extra TFIIIC (ETC) without RNAPIII, are not bound by Fpt1. Therefore, TFIIIC alone cannot explain differential binding of Fpt1 at tRNA genes. Moreover, in the tDNA metagene analysis, some ChIP-exo peaks of Fpt1 are not explained by RNAPIII, TFIIIB, and TFIIIC (left-most Fpt1 peak in **Figure S2C-H**). This may point to direct contacts between Fpt1 and DNA sequences upstream of the tDNA gene bodies.

Taken together, our data extend the current knowledge on regulation of tDNAs by RNAPIII and provide evidence for a chromatin-associated regulatory mechanism embedded in the tDNA proteome (as outlined in **Figure 7**). In conditions of active transcription, Maf1 is phosphorylated and located in the cytoplasm. In the nucleus, the chromatin-bound fraction is composed of the RNAPIII transcription machinery that facilitates tDNA transcription. At the tRNA gene, RNAPIII and TFIIIB are strongly enriched while TFIIIC and Fpt1 are moderately enriched. In repressive conditions, Maf1 is dephosphorylated and imported into the nucleus where it directly interacts with RNAPIII and TFIIIB in the unbound fraction. This prevents assembly of TFIIIB and RNAPIII onto the tDNA. In the chromatin-bound fraction, occupancy of TFIIIB and RNAPIII is decreased while TFIIIC and Fpt1 show increased occupancy at the tRNA gene, supporting a model of competition. In repressive conditions, even though Maf1 prevents RNAPIII assembly and is the major repressor, Fpt1 is required to promote RNAPIII eviction and loading of TFIIIC. Fpt1 may do so by acting on RNAPIII via TFIIIB, or by stabilizing TFIIIC. The latter would be in line with the ChIP-exo correlations between Fpt1 and TFIIIC and the proposed dual role of TFIIIC, promoting activation as well as repression of tDNA transcription [52, 53]. The mechanism by which Fpt1 acts on TFIIIB and TFIIIC remains to be explored.

**Figure 7.**
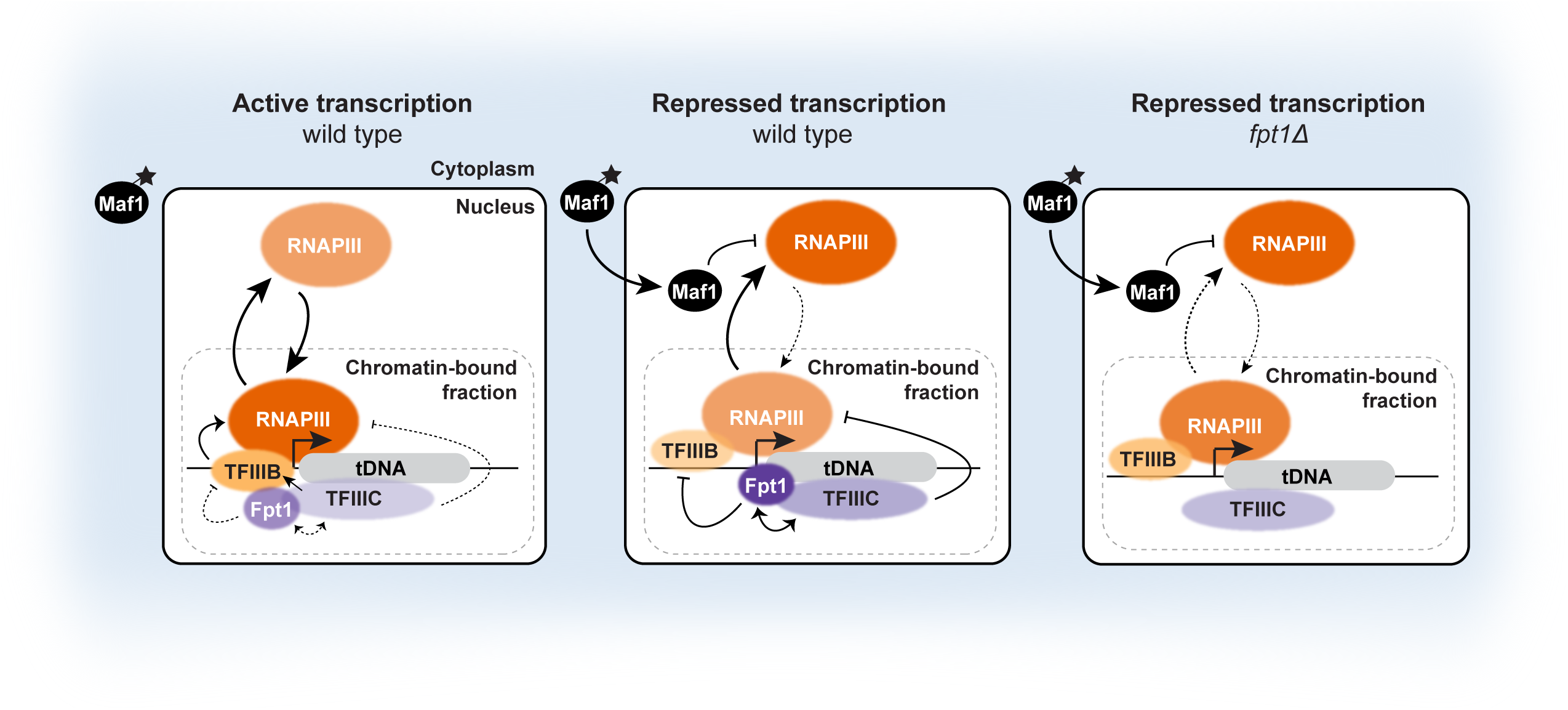
A model for chromatin-associated regulation of the tDNA transcription machinery. In conditions of active transcription, Maf1 is phosphorylated and located in the cytoplasm (shown in blue). In the chromatin-bound fraction in the nucleus, RNAPIII and TFIIIB show high occupancy while TFIIIC and Fpt1 show low occupancy. TFIIIC promotes binding of TFIIIB, which in turn promotes recruitment of RNAPIII and there is active exchange of RNAPIII molecules between the unbound and bound fraction (indicated with arrows). In repressive conditions, Maf1 is dephosphorylated and imported into the nucleus where it directly interacts with RNAPIII and TFIIIB in the unbound fraction. This prevents assembly of TFIIIB and RNAPIII onto the tDNA, resulting in decreased recruitment of RNAPIII molecules. In the chromatin-bound fraction, occupancy of TFIIIB and RNAPIII is decreased while TFIIIC and Fpt1 show increased occupancy at the tRNA gene, supporting a model of competition between the core tDNA transcription complexes. The behavior of Fpt1 and the changes that happen in its absence support a model in which, in repressive conditions (and to a lesser extent in active conditions), Fpt1 promotes RNAPIII eviction and loading of TFIIIC either via de-stabilization of TFIIIB or by stabilizing TFIIIC. In the absence of Fpt1, TFIIIC occupancy is reduced and RNAPIII and TFIIIB are partially retained at the chromatin, causing improper adjustment of the tDNA proteome to changing nutrient conditions.

Understanding how Fpt1 interplays with the tDNA proteome and how it affects RNAPIII dynamics and cellular fitness will provide insights into how yeast as a workhorse in biotechnology can be metabolically optimized for production of biomolecules. Phylogenetic analysis indicates that the *FPT1* gene evolved in the Saccharomycetales clade closely related to *Saccharomyces cerevisiae.* Fpt1 likely has a yeast-specific origin and based on the protein sequence, no Fpt1 homologues have been identified in multicellular eukaryotes. However, we expect that chromatin-associated regulatory mechanisms of RNAPIII assembly are present in other eukaryotes as well. Current advances in genome engineering and proteomics may facilitate the development of strategies to decode tDNAs by barcode sequencing or capture approaches in more complex eukaryotes to explore mechanisms of chromatin-associated regulation of RNAPIII. Our study emphasizes the importance of not overlooking uncharacterized proteins in such efforts, as they may possess novel regulatory roles that could change our views on fundamental cellular processes [65].

## MATERIAL AND METHODS

### Yeast strains, oligos and plasmids

All yeast strains, oligos and plasmids used in this study are listed in the Supplemental Material & Methods. Yeast strains were maintained in YEPD at 30°C unless otherwise specified. Details on growth conditions, media compositions, and strain and plasmid construction are provided in the Supplemental Material & Methods. *YKR011C* is referred to as *FPT1* (Factor in the Proteome of tDNAs number 1). This name has been reserved in the Yeast Genome Database: www.yeastgenome.org/locus/S000001719.

### Chromatin immunoprecipitation

Chromatin immunoprecipitation was performed as previously described in [43] with the following modifications. Cell cultures were grown in 75 mL YEPD until mid-log phase (approximately 0.5-1*10^7^ cells/mL, OD_660_ 0.5-0.8) and cross-linked for 10 minutes (Rpo31-TAP ChIP) or 15 minutes (Epi-Decoder, ChIP-sequencing, Fpt1-TAP ChIP) with one-tenth of freshly prepared fix solution (11% formaldehyde, 50 mM Hepes-KOH [pH 7.5], 100 mM NaCl, 1 mM EDTA). Cross-linking was quenched for 5 minutes with glycine (125 mM final concentration) or 1 minute with Tris-HCl pH = 8.0 (750 mM final concentration). Chromatin shearing was performed using the Bioruptor PICO (Diagenode) for 6-10 minutes (depending on the experiment) with 30-second intervals at 4°C. For chromatin immunoprecipitation, one volume of chromatin was mixed with 1/10 volume of Dynabeads M-270 Epoxy (ThermoFisher, LOT 01063817) coupled to immunoglobulin G (IgG) and incubated overnight on a turning wheel at 4°C.

### Epi-Decoder

Epi-Decoder was performed as described in [43]. Briefly, a NATMX cassette was inserted in proximity to the tDNA-Ty1 locus. Subsequently, a gRNA-containing CRISPR-Cas9 plasmid was used to integrate a random 16 bp DNA-barcode oligo library in between the divergent tRNA and Ty1 genes. Colonies were picked and arrayed in a 384-format library to construct a barcoded yeast library. The barcoded yeast library was crossed with a TAP-tag protein library using SGA to create an Epi-Decoder library. Colonies on 384-format Epi-Decoder library plates were pooled together and grown to mid-log phase. Three different barcode-protein combinations represent biological replicates. Just before fixation, the biological replicates were split in three technical replicates (that have the same barcode-protein combinations). Details on sample preparation and data analysis are described in the Supplemental Material & Methods.

### ChIP-sequencing

DNA from chromatin immunoprecipitation was prepared for sequencing with the KAPA Hyper Prep Kit (KAPA Biosystems). DNA was amplified with 19 (IP) or 11 cycles (input) and a double sized 0.6X and 1X selection was performed to select for 200-450 bp DNA fragments using AMPure XP beads (Beckman Coultier). An additional 1X size selection was performed on samples that showed high primer-dimer peaks. DNA concentration was measured using a Qubit dsDNA HS Assay Kit (Invitrogen) and size-distribution of DNA fragments was visualized with an Agilent DNA 1000 Kit (Agilent Technologies) and Agilent 2100 Bioanalyzer. Purified DNA was sequenced (single read, 65 bp) on a HiSeq2500 platform (Illumina). Data analysis is described in the Supplemental Material & Methods.

### ChIP-exo

All yeast cultures were grown as described previously [66]. Briefly, cultures were grown at 25°C in 50 mL YEPD medium to an OD_600_ of 0.6-0.8. Cells were cross-linked at room temperature with formaldehyde (1% v/v, 15 min) and quenched by glycine (125 mM, 5 min). Cross-linked cells were centrifuged at 4°C (4000 rpm, 5 min) and washed with chilled ST buffer (10 mM Tris-HCl, pH7.5 and 100 mM NaCl). Cell pellets were frozen and stored at -80°C until further use. Cells were lysed and isolated chromatin was sheared as described previously [66]. For ChIP-exo 5.0, steps were performed as described previously [67]. A single ChIP experiment used chromatin obtained from 50 mL culture and 40 µL IgG-Dynabeads slurry. ChIP-exo libraries were verified on agarose gels and purified. Subsequent steps for library sequencing, quality control, and data analysis (including NCIS normalization) were performed as described previously [66].

### ChIP-qPCR

Quantitative PCR was performed on a LightCycler 480 II (Roche) and analyzed with software from the manufacturer. In total, 4.2 µL purified DNA from chromatin immunoprecipitation was mixed with 5 µL 2x SensiFAST SYBR No-ROX kit (Bioline) and 0.4 µL forward and reverse primer (10 µM). The quantity of original DNA was determined by interpolating the resulting Cp values from a linear standard curve of values obtained from the dilution-series.

### Live cell imaging

Live-cell imaging was performed as previously described in detail [68] with minor modifications. In brief, cells were grown to OD_600_ 0.2-0.4 in SC + 2% glucose. For repressive conditions, cells were subsequently subjected to SC + 2% ethanol (at 30°C) or SC + 2% glycerol (at 37°C) for 2 hours. Cells were imaged on a coverslip with an agarose pad consisting of 2% agarose in SC + 2% glucose, ethanol or glycerol at 30°C. Microscope settings and data analysis are described in the Supplemental Material & Methods.

### Growth assays and flow cytometry

Growth on solid media was measured using spot test analysis. Serial ten-fold dilutions were spotted on plates containing SC media including 2% glucose, glycerol or ethanol. Growth was assessed after 2 days at 30°C or 37°C. For competitive growth assays in liquid media, cells were cultured 48h in pre-competition media (SC + 2% glucose). Wild-type and *fpt1*Δ cells (0.5*10^6^), labelled with mScarlet and NeonGreen fluorescent markers, were mixed in 1 mL SC + 2% glucose (t = 0). A color swap was done to account for reporter effects on cell fitness. Additionally, NeonGreen labelled wild type and mScarlet labelled wild type were mixed to measure fitness defects caused by the fluorescent reporters. Culture conditions for the competitive growth assay and flow cytometry analysis are described in the Supplemental Material & Methods.

### RNA-sequencing

Cells were grown in 10 mL YEPD to mid-log phase. Half of the culture was spun down (2000 rpm, 5 minutes) and media was changed to YEP + 2% ethanol. Cells were grown for another 2h. Pellets were harvested by a 2 minute spin at 3000 rpm 4°C, washed once with water and stored at -80°C. RNA was isolated using the Qiagen RNeasy kit according to the manufacturer’s protocol. DNase I treatment was performed on the column (NEB M0303S) and RNA was stored at -80°C. The Illumina TruSeq Stranded mRNA kit was used to make cDNA library by following the manufacturer’s protocol. cDNA was sequenced (single read, 65 bp) on a HiSeq2500 platform (Illumina). Reads were mapped to SacCer3 using STAR (version 2.7.1a) [69] and count tables were generated using the bioconda package eXpress [70]. Differential expression analysis was performed with DEseq2 [71]. Gene ontology analysis on differentially expressed genes was performed using ShinyGO 0.77 [72].

## AUTHOR CONTRIBUTIONS

M.E.v.B. performed and analyzed the experiments and wrote the paper. I.v.K. constructed the tDNA-Ty1 Epi-Decoder library, performed the initial tDNA-Ty1 Epi-Decoder analysis in glucose and performed and analyzed the RNA-sequencing experiment. C.M. performed and analyzed the ChIP-exo experiments. C.L. and R.J.C.K. analyzed Epi-Decoder sequencing data. I.B. performed and analyzed microscopy experiments. T.v.d.B. analyzed ChIP-seq data. R.M. helped with discussing the methods used for analyzing the Epi-Decoder data. T.v.W. helped with constructing the tDNA-Ty1 Epi-Decoder library. A.D.C. and K.V. performed the phylogenetic analysis. M.M. helped with flow cytometry data acquisition and analysis. T.L.L. and R.B. helped with discussing the project and supervision. B.F.P. helped with ChIP-exo data analysis and supervision. F.v.L. conceived and supervised the project, analyzed data, and wrote the paper.

## ACKNOWLEDGEMENTS

We thank David Tollervey and Tomasz Turowksi for sharing the CRAC dataset on RNAPIII transcripts and discussing the project. We thank Evelina Tuttuci for sharing the pET542 and pET543 plasmids and Johan van Heerden and Frank Bruggeman for discussing the project. We thank Ben Morris for help with re-arraying yeast libraries by robotics and Pascale Daran-Lapujade for advice regarding the neutral X-2 locus. We thank the Genomics Core Facility, Robotics Facility, High Throughput Screening Facility, and Flow Cytometry Facility of the NKI for assistance. We thank members of the F.v.L. lab for helpful discussions. We acknowledge the following resources: ScriptManager, RRID:SCR_021797; Platform for Epigenomic and Genomic Research, RRID:SCR_021861; Galaxy, RRID:SCR_006281; Picard, RRID:SCR_006525; Cornell University Biotechnology Resource Center Epigenomics Core Facility, RRID:SCR_021287; Cornell University BRC Genomics Core Facility, RRID:SCR_021727; Pennsylvania State University’s Institute for Computational and Data Sciences Advanced Cyberinfrastructure.

## DATA AVAILABILITY

Sequencing data for ChIP-seq (including bigwig files), RNA-seq (including FKPM count data) and ChIP-exo (including BedGraphs files) are available from the GEO database with accession number GSE227470. De-multiplexed Epi-Decoder sequencing data and TAP-barcode combinations from Epi-Decoder libraries used in this study have been submitted to the NCBI BioProject database under accession number PRJNA945378. All processed data are within the paper and the Supplemental Material & Methods. Previously published ChIP-exo datasets used in this study can be accessed from the GEO database with accession number GSE147927 [58]. Epi-Decoder data from the HO locus can be found at the NCBI BioProject database under accession number PRJNA610036 (SRX7842936) [41]. Additional details about data analysis, custom scripts and parameters used for ChIP-exo are available on Github (https://github.com/CEGRcode/2023-Breugel_JournalXXXX).

