## Supplemental Material & Methods for "A chromatin-associated regulator of RNA Polymerase III assembly at tRNA genes revealed by locus-specific proteomics"

### SUPPLEMENTAL MATERIAL AND METHODS

#### Supplemental Figure S1

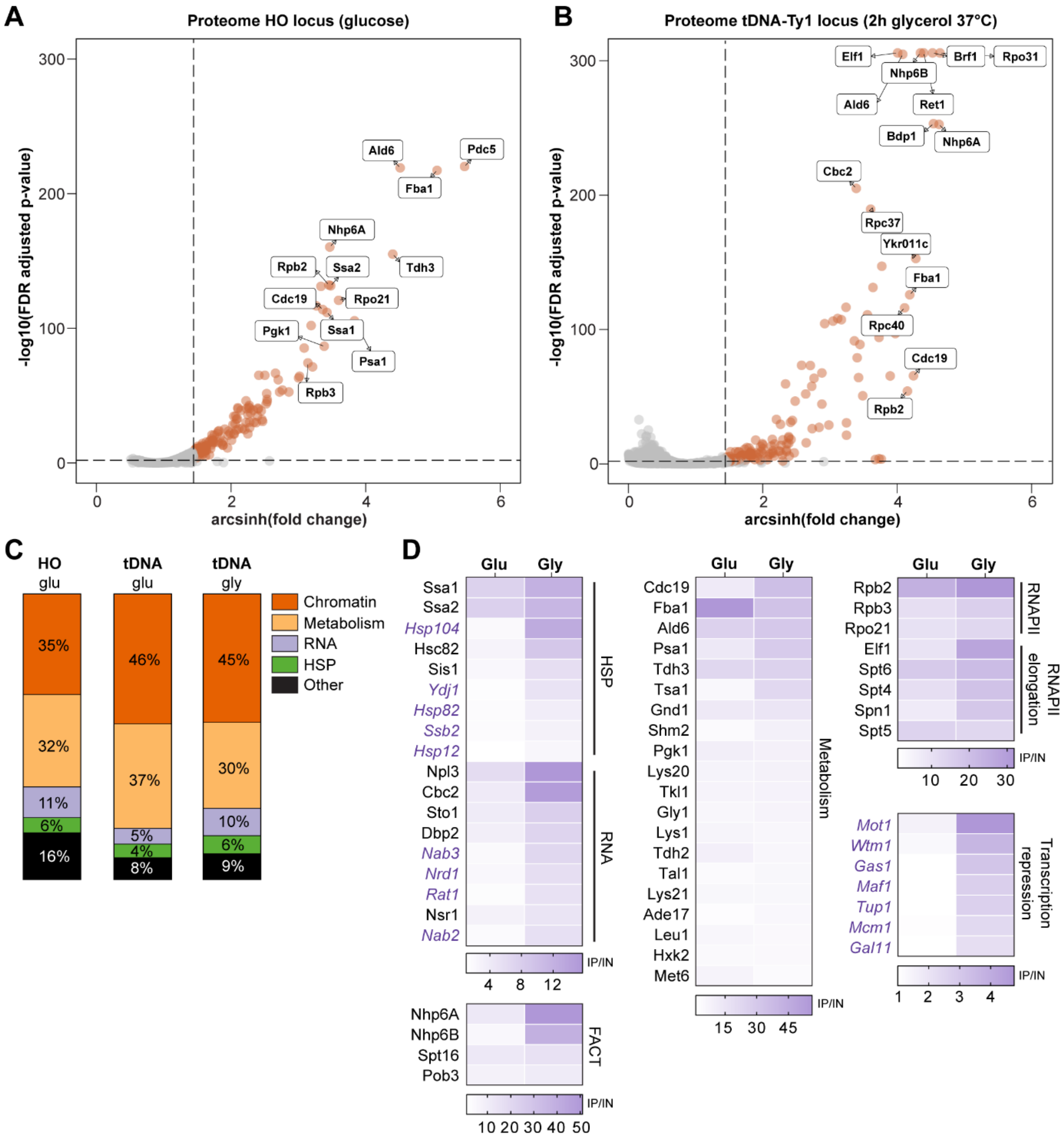


**Figure S1. The proteome of a tDNA locus is rewired in response to nutrient availability. (A)** Fold change between ChIP and input (arcsinh) and FDR (p-value) for 3705 proteins at the HO locus in glucose (data from [1]). Data describes three biological replicates (different DNA-barcodes) and three technical replicates (same DNA-barcode). Proteins classified as 'binder' are indicated with colored dots and significance thresholds with dashed lines (basemean ≥ 400, FDR ≤ 0.01 and $\log_2$ fold change ≥ 1). **(B)** As in (A), fold change between ChIP and input (arcsinh) and FDR (p-value) for 3681 proteins at the tDNA-Ty1 locus in glycerol 2h (37°C). **(C)** Fraction of proteins classified as 'binder' in glucose and glycerol 2h (37°C) at the HO or tDNA-Ty1 locus for different functional classes: chromatin, metabolism, RNA, heat shock proteins (HSP) and others. **(D)** Heat maps show fold change values (ChIP vs input) for different protein complexes or families at the tDNA-Ty1 locus. The average ChIP/input of three biological replicates is shown. Protein names are color-coded based on the Euler diagram shown in Figure 1D. Black: 'binder' in glucose and glycerol 2h (37°C). Purple and italic: 'binder' in glycerol 2h (37°C).

E

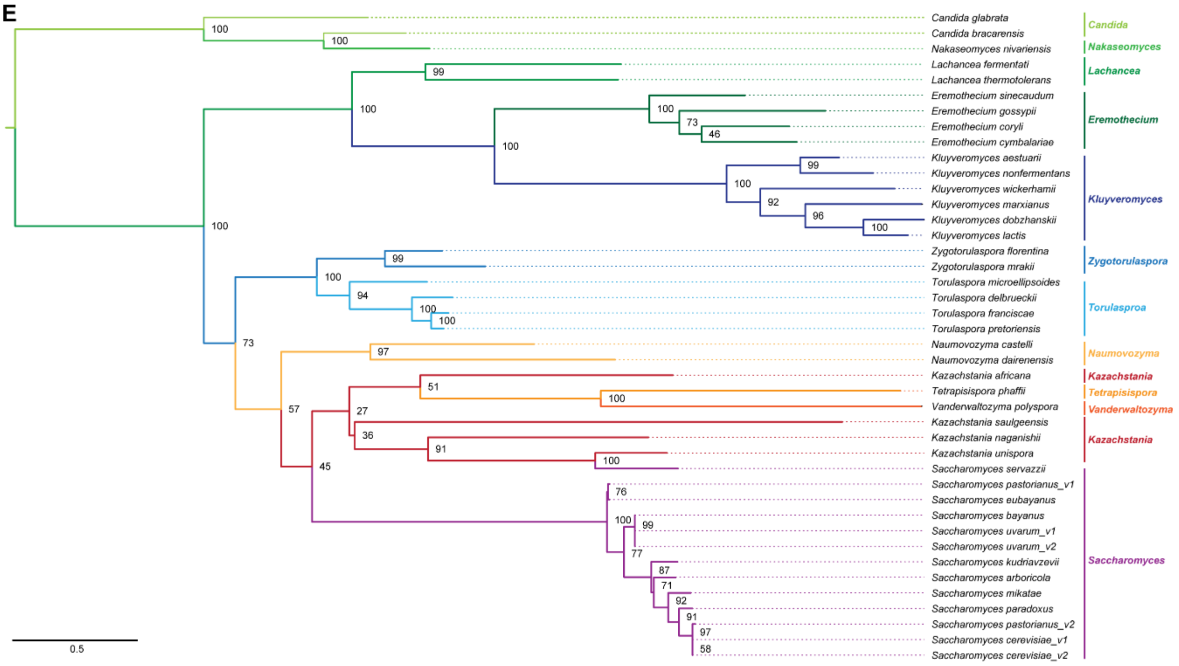


**Figure S1E. The proteome of a tDNA locus is rewired in response to nutrient availability. (E)** Maximum Likelihood phylogenetic tree inferred from amino acid alignment of 42 Ykr011c orthologues using IQ-tree. Bootstrap values are shown on the nodes. The scale represents 0.5 substitutions per amino acid position.

### Supplemental Figure S2

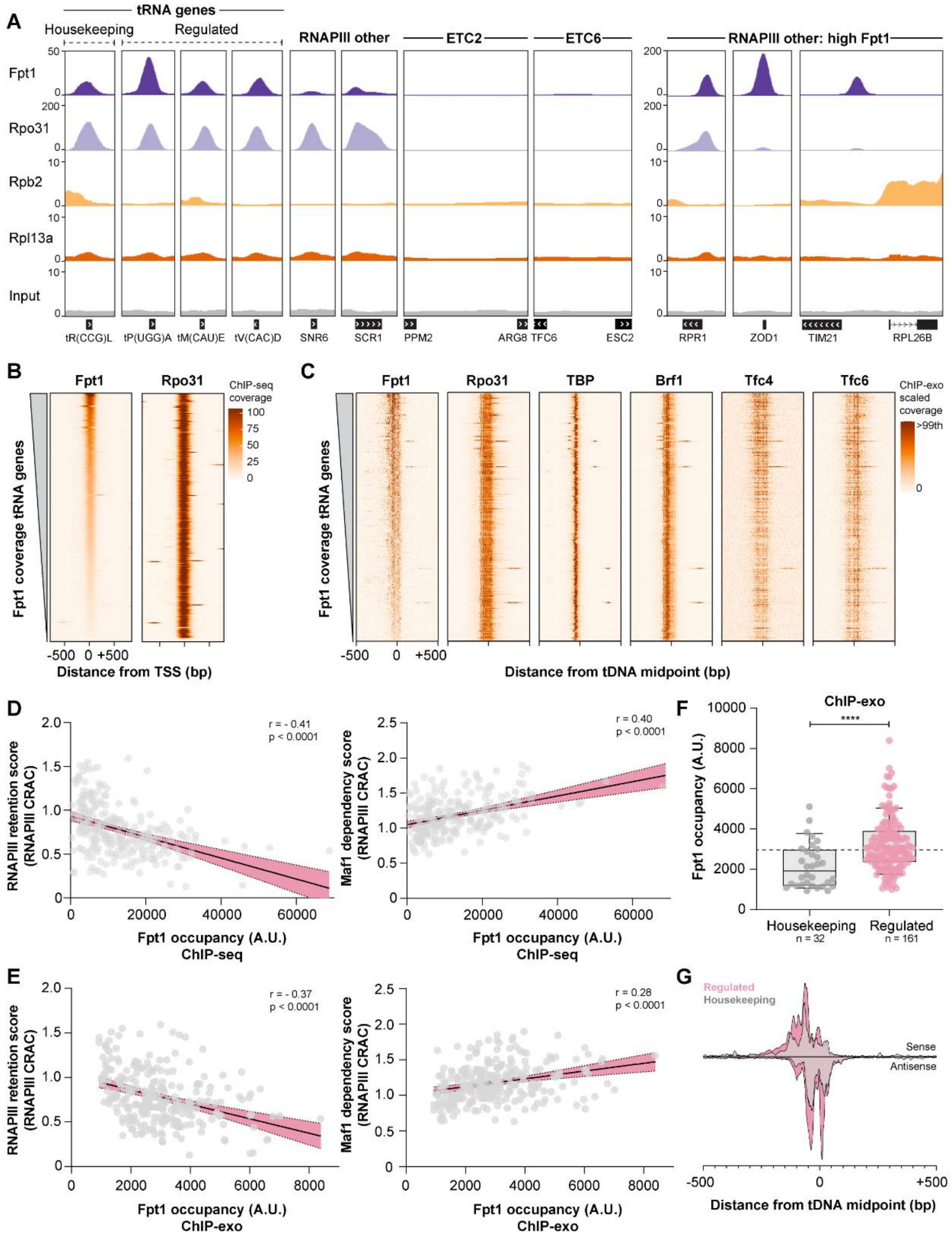

**Figure S2. Fpt1 occupancy at tDNAs is variable and correlates with the response to nutrient perturbations. (A)** Genome tracks show pooled ChIP-seq data of three biological replicates for TAP-tagged Fpt1, Rpo31, Rpb2 and Rpl13a at tRNA genes *tR(CCG)L*, *tP(UGG)A*, *tM(CAU)E* and *tV(CAC)D*, RNAPIII transcribed genes *SCR1* and *RPR1*, ETC sites *ETC2* and *ETC6* and tDNA relics *ZOD1* and *iYGR033c* (located between *TIM21* and *RPL26B*). **(B)** Fpt1 and Rpo31 coverage at tRNA genes (n = 275) from ChIP-seq data ranked from high to low Fpt1 occupancy. **(C)** Fpt1, Rpo31, TBP, Brf1, Tfc4 and Tfc6 coverage (up to the 99th percentile) at tRNA genes (n = 261) from ChIP-exo data ranked from high to low Fpt1 occupancy. **(D-E)** Pearson correlation plots of RNAPIII retention scores and Maf1 dependency scores from [2] with Fpt1 occupancy from ChIP-seq at 243 tRNA genes (in D) and from ChIP-exo at 249 tRNA genes (in E). Pearson coefficient and p-value (two-tailed) are indicated in the top right corner. **(F)** Fpt1 occupancy (ChIP-exo) at housekeeping tRNA genes (n = 32) and regulated tRNA genes (n = 161). The average Fpt1 occupancy (2983) across 249 tRNA genes is indicated with a dotted horizontal line. Significance was determined using a Welch-corrected unpaired two-tailed t-test, ****: $p < 0.0001$. **(G)** Fpt1 ChIP-exo metagene plot of housekeeping (n = 32) and regulated (n = 161) tRNA genes.

### Supplemental Figure S3

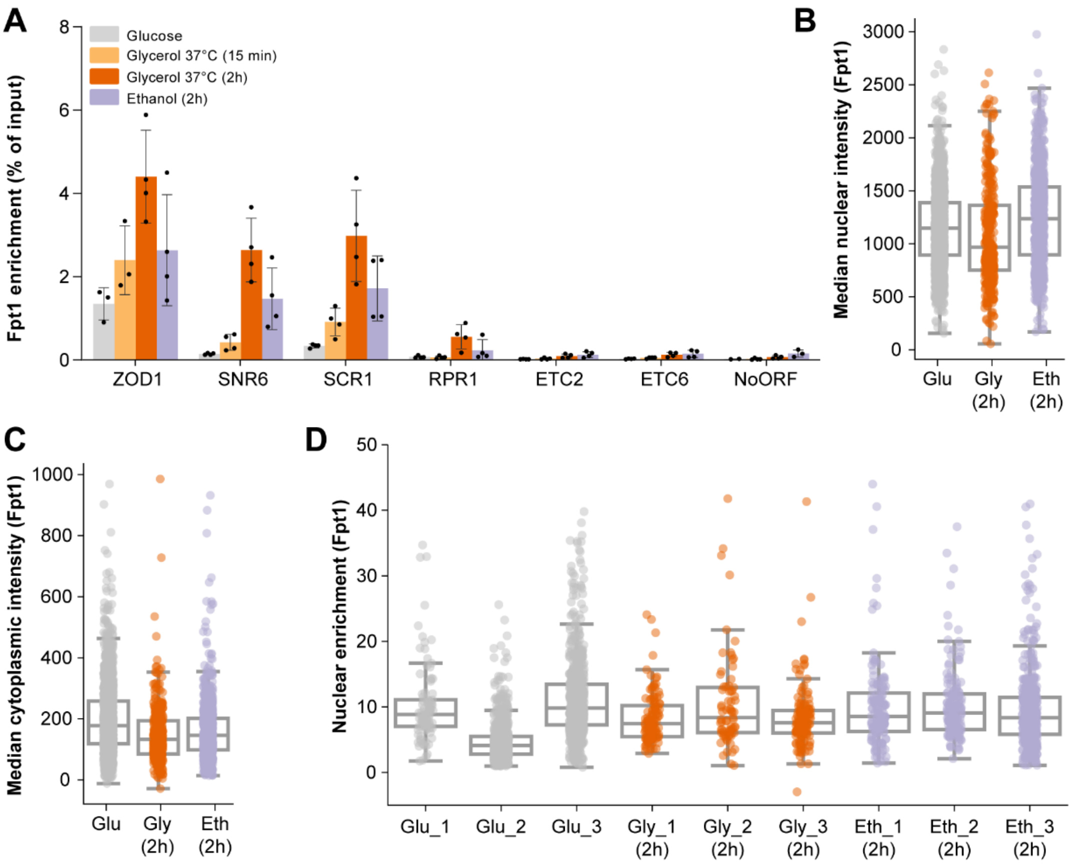


**Figure S3. Fpt1 occupancy and nuclear localization in active and repressive conditions. (A)** Fpt1 enrichment (ChIP) at the tDNA relic *ZOD1*, the RNAPIII transcribed non-tRNA genes *SNR6*, *SCR1* and *RPR1*, the ETC sites *ETC2* and *ETC6* and a negative control locus (NoORF). The average and standard deviation of four biological replicates is shown (unless a replicate was excluded from the qPCR data due to technical variation). **(B)** Median nuclear intensity in three biological replicates of Fpt1 in glucose (n = 1536 cells), glycerol 2h 37°C (n = 327 cells) and ethanol 2h (n = 771 cells). Circles show data for individual cells and box plots show the distribution of the data where the box indicates the quartiles and whiskers extend to show the distribution, except for outliers. **(C)** As in (B), showing cytoplasmic intensity of Fpt1. **(D)** Nuclear enrichment of Fpt1 as shown in Figure 3D for three individual biological replicates.

### Supplemental Figure S4

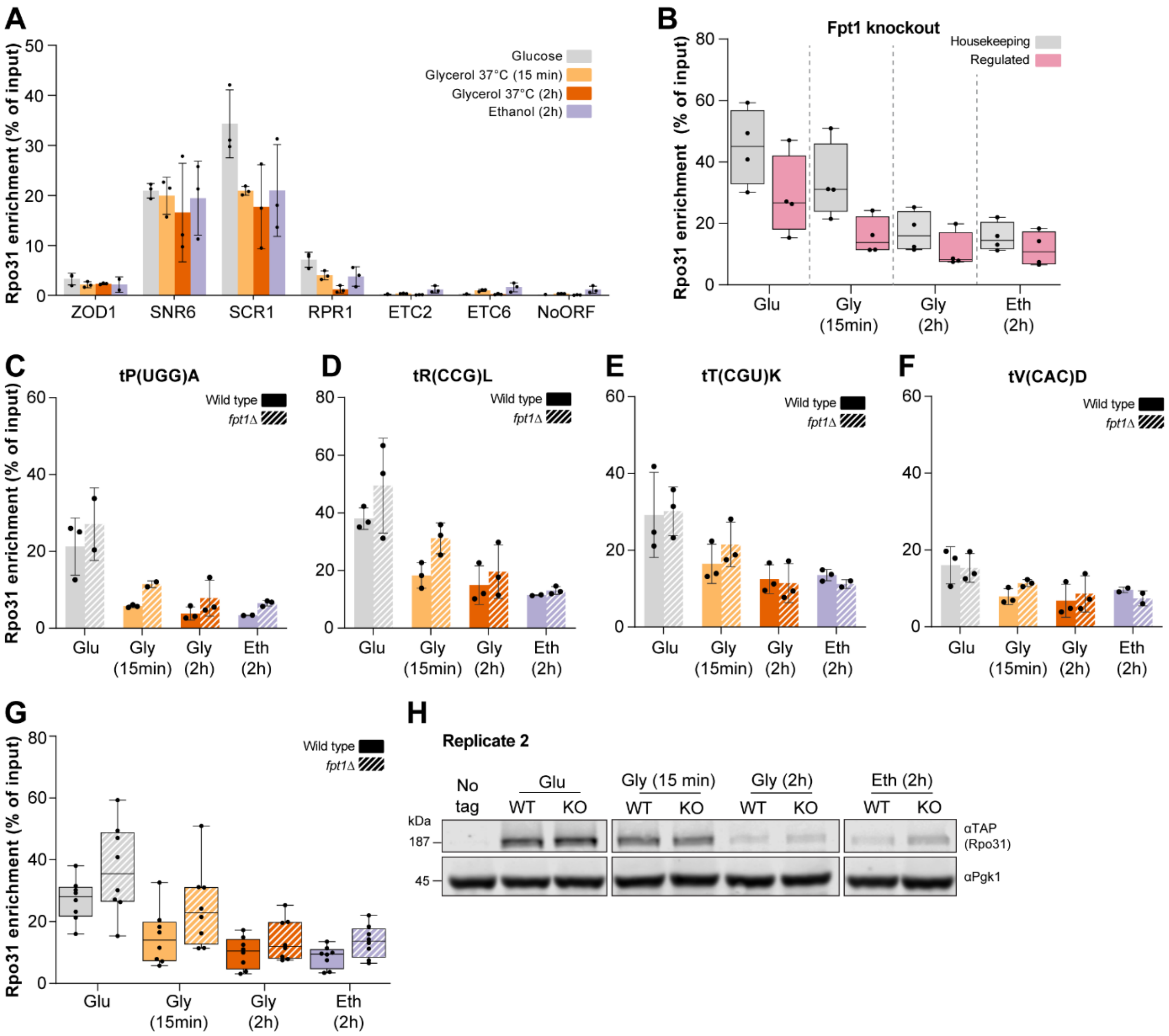


**Figure S4. Fpt1 controls the dynamics of RNAPIII upon changing nutrient conditions. (A)** Rpo31 enrichment (ChIP) at the tDNA relic *ZOD1*, the RNAPIII transcribed non-tRNA genes *SNR6*, *SCR1* and *RPR1*, the ETC sites *ETC2* and *ETC6* and a negative control locus (NoORF). The average and standard deviation of three biological replicates is shown (unless a replicate was excluded from the qPCR data due to technical variation). **(B)** Rpo31 enrichment (ChIP) in *fpt1Δ* at housekeeping tRNA genes (grey, n = 4) and regulated tRNA genes (pink, n = 4) in glucose, glycerol 2h or 15 min (37°C) and ethanol 2h. For each tRNA gene, the average of three biological replicates is shown. Whiskers indicate the minimum and maximum. **(C-F)** Rpo31 enrichment (ChIP) in wild type and *fpt1Δ* at different tRNA genes in glucose, glycerol 2h and 15 min (37°C), and ethanol 2h. The average and standard deviation of three biological replicates is shown (unless a replicate was excluded from the qPCR data due to technical variation). **(G)** Rpo31 enrichment (ChIP) at eight tRNA genes (as shown in Figure 4C-F and S4C-F) in wild type and *fpt1Δ*. For each tRNA gene, the average of three biological replicates is shown. Whiskers indicate the minimum and maximum. **(H)** Biological replicate 2 of Rpo31-TAP (anti-TAP) immunoblot in glucose, glycerol 2h and 15 min (37°C), and ethanol 2h. WT: wild type, KO: *fpt1Δ*. Pgk1 was used as a loading control. No tag is the negative control strain lacking a TAP-tag (BY4741).

### Supplemental Figure S5

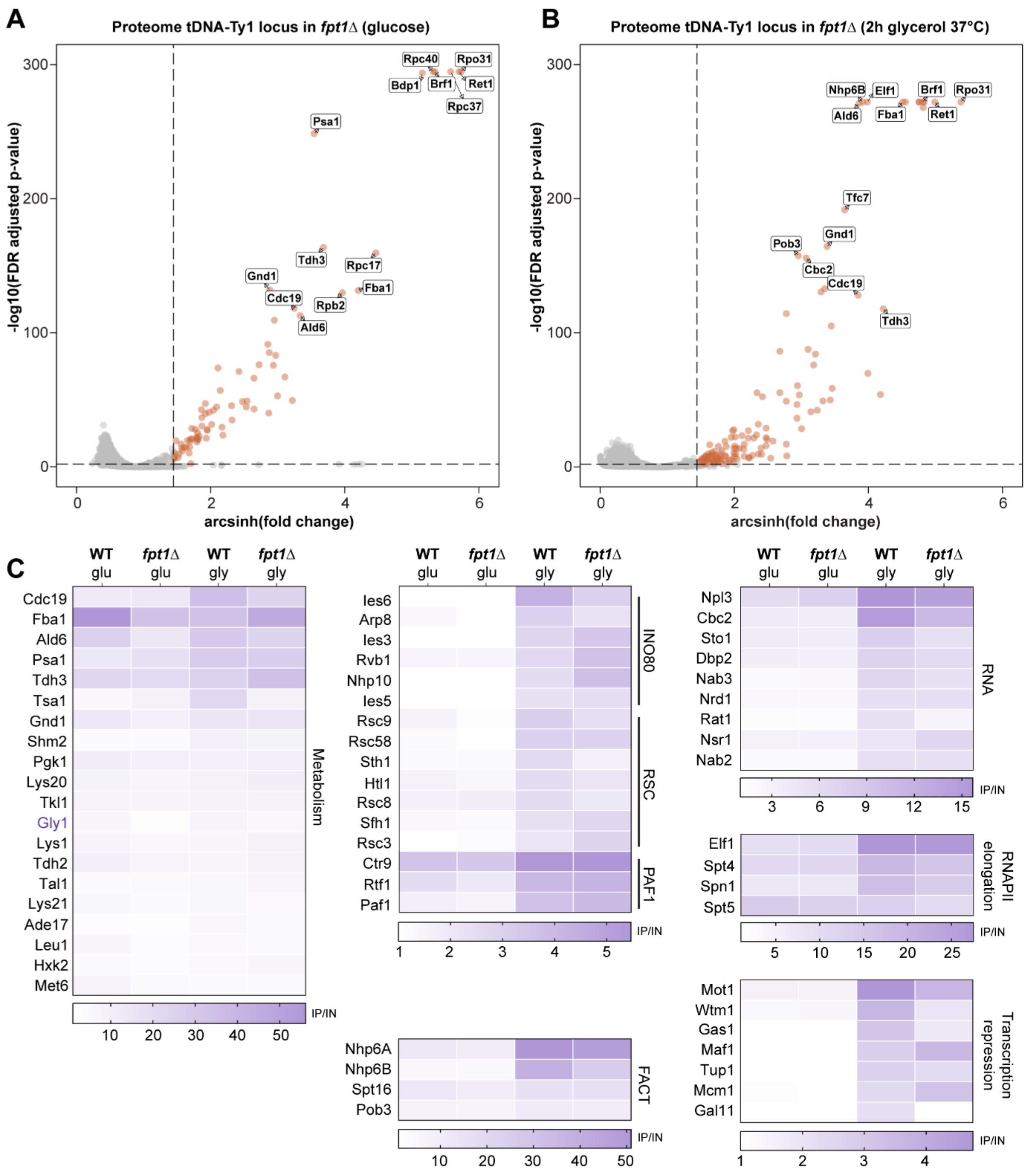


**Figure S5. Fpt1 causes reprogramming of the RNAPIII transcription machinery. (A)** Protein occupancy score in *fpt1Δ* at the tDNA-Ty1 locus in glucose for 3707 proteins: fold change between ChIP and input (arcsinh) and FDR (p-value). Data describes three biological replicates (different DNA-barcodes) and three technical replicates (same DNA-barcode). Proteins classified as 'binder' are indicated with colored dots and dashed lines show significance thresholds (basemean ≥ 400, FDR ≤ 0.01 and $log_2$ fold change ≥ 1). **(B)** As in (A), fold change between ChIP and input (arcsinh) and FDR (p-value) for 3673 proteins in *fpt1Δ* at the tDNA-Ty1 locus in 2h glycerol (37°C). **(C)** Heat maps show fold change values (ChIP vs input) at the tDNA-Ty1 locus for different protein complexes or families for wild type or *fpt1Δ* in glucose and 2h glycerol (37°C). WT = wild type, KO = *fpt1Δ*, glu = glucose, gly = glycerol 2h (37°C). The average ChIP/input of three biological replicates is shown.

### Supplemental Figure S6

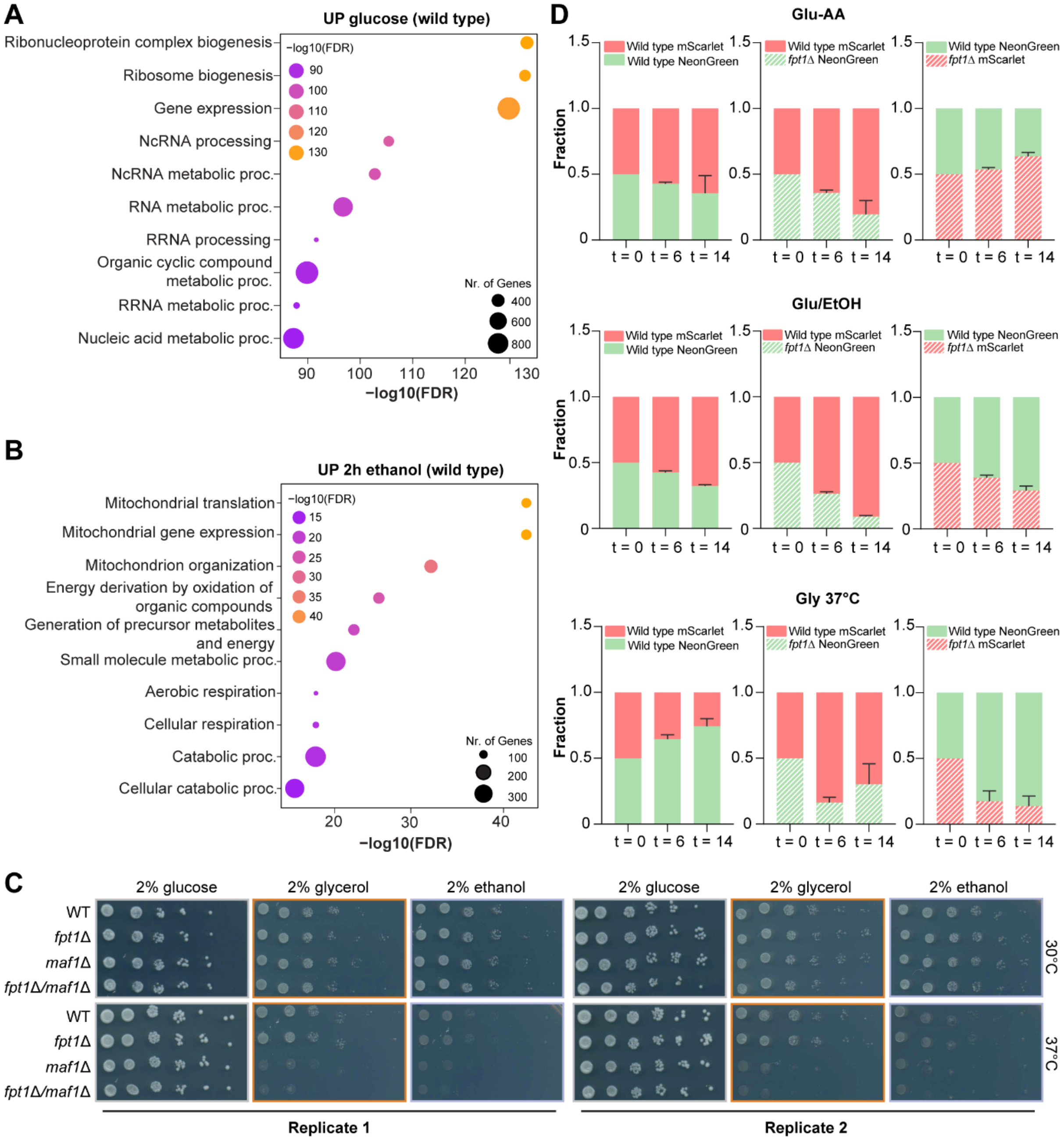


**Figure S6. Loss of Fpt1 alters shutdown of ribosome biogenesis gene expression and cellular fitness in repressive conditions. (A)** Gene ontology analysis of genes upregulated in wild type in glucose (n = 2026). **(B)** Gene ontology analysis of genes upregulated in wild type in ethanol 2h (n = 1929). **(C)** Spot test analysis of different mutant backgrounds in SC + 2% glucose, glycerol and ethanol for two biological replicates at 30 °C and 37 °C. In repressive conditions, *maf1Δ* cells showed temperature sensitivity, which has been partly attributed to the loss of expression of two gluconeogenic enzymes [3]. **(D)** The ratio of red and green cells at three different time points (t = 0, t = 6 or t = 14 days) of a competitive growth assay (see Figure 6E-F). For each growth condition, three combinations were tested: wild type red + wild type green, wild type red + *fpt1Δ* green and wild type green + *fpt1Δ* red. Bars indicate the mean value of four biological replicates and the error bars indicate the standard deviation.

**Strain and plasmid construction**

To construct strains (**Supplemental Table S9**), *S. cerevisiae* was transformed using the LiAc/ssDNA method as described in [4]. For standard transformations, BY4741 was used as a parental strain, unless otherwise specified.

Epi-Decoder library manipulations on solid media were performed using synthetic genetic array (SGA) technology [5] and a ROTOR instrument (Singer Instruments, Watchet, UK). The NKI4214 TAP-tag library was made by crossing NKI4212 with the commercially available yeast TAP-tagged ORF library. Strains NKI5623, NKI4221, NKI5625 and NKI5626 were picked from the NKI4214 library. NKI5601 was made by using plasmid pFvL099 (**Supplemental Table S10**) and the *NatMX Epi-Decoder locus* oligos (**Supplemental Table S11**). The NKI5614 barcode library was made by transforming NKI5601 with the CRISPR-Cas9 plasmid pIla011, containing a gRNA targeting the tDNA-Ty1 locus, and a PCR amplified barcode oligo library (**Supplemental Table S11**) acting as repair template. The NKI5616 Epi-Decoder library was constructed by crossing the NKI4214 and NKI5614 libraries. Strains NKI5662-NKI5667 were picked from the NKI5616 Epi-Decoder library. NKI5633 was constructed using the pRS400 plasmid and *fpt1Δ* oligos. The NKI5642 *fpt1Δ* Epi-Decoder library was made by crossing NKI5616 with NKI5633.

NKI5620 was isolated from the YSC1053 knockout library from Open Biosystems. NKI5632 was created using plasmid pFvL029 and *Fpt1-TAP* oligos. NKI2580 and NKI2581 were constructed using plasmid pFvL099 and *maf1Δ* oligos. NKI2581 originated from NKI5633. NKI5753 was made using the pKT127 plasmid and *Fpt1-GFP* oligos. NKI5756 originated from NKI5753, which was transformed with Pac1 digested pTL306 plasmid to insert a fluorescent coat protein and URA3 selection marker at the *URA3* locus. NKI5707 and NKI5724 were constructed by amplifying NeonGreen from pET542 with the *NeonGreen/Scarlet X-2 locus* oligos and using the CRISPR-Cas9 containing pMvB03 plasmid targeting the X-2 locus (a neutral intergenic locus located at ChrX:195625-195645 as described in [6]). NKI5712 and NKI5726 were constructed similarly but instead the mScarlet containing pET543 plasmid was used.

Plasmid pMvB03 was constructed by cloning the X-2 gRNA into pML104-NatMX3 using the BclI and SwaI restriction sites. Plasmid pIla011 was constructed by cloning the tDNA-Ty1 gRNA into pML104-URA3 using the BclI and SwaI restriction sites.

**Growth conditions and media compositions**

For all experiments (except the competitive growth assay and ChIP-exo), liquid yeast cultures were grown to mid-log phase in 2% glucose (YEPD) at 30°C. Optionally, cells were switched to repressive conditions by incubating for 2 hours in 2% ethanol (YEPEtOH) at 30°C or 2% glycerol (YEPgly) at 37°C. For ChIP-exo, cells were grown in 2% glucose (YEPD) at 25°C. In the competitive growth assay, cultures were grown in synthetic complete (SC) media using the following recipes:

**Glu-AA:** Switching back and forth between SC + 2% glucose (AA complete) and yeast nitrogen base (YNB) + auxotrophic AA + 2% glucose. *SC + 2% glucose***:** 6.7 g YNB w/o amino acids, carbohydrates and with ammonium sulfate + 2 g drop-out mix complete w/o YNB + 100 mL 20% glucose + 900 mL Milli-Q. *YNB + auxotrophic AA + 2% glucose*: 100 mL 10X YNB (with ammonium sulfate, w/o amino acids) + 200 mL 250 mM potassium hydrogen phthalate + 10 mL methionine (5 mg/mL) + 10 mL histidine (5 mg/mL) + 10 mL leucine (10 mg/mL) + 100 mL uracil (1 mg/mL) + 10 mL adenine (1 mg/mL) + 10 mL myo-inositol (100X solution of 0.2 mg/mL) + 10 mL 4-amino benzoic acid potassium salt (100X solution of 0.02 mg/mL) + 100 mL 20% glucose + 440 mL Milli-Q.

**Gly 37°C:** Continuous growth in SC + 2% glycerol. *SC + 2% glycerol*: 6.7 g YNB w/o amino acids, carbohydrates and with ammonium sulfate + 2 g drop-out mix complete w/o YNB + 100 mL 20% glycerol + 900 mL Milli-Q.

**Glu/EtOH:** Switching back and forth between SC + 2% glucose and SC + 2% ethanol. *SC + 2% ethanol*: 6.7 g YNB w/o amino acids, carbohydrates and with ammonium sulfate + 2 g drop-out mix complete w/o YNB + 20 mL 100% ethanol + 980 mL Milli-Q.

Media compositions for solid agarose plates used in Epi-Decoder are described in [7]. Solid agarose plates used in spot test analysis (**Figure S6C**) contained synthetic complete (SC) media including 2% glucose, ethanol or glycerol.

**Epi-Decoder sample preparation and data analysis**

Chromatin samples were sonicated for 10 minutes with 30 second intervals. To prepare libraries for sequencing, barcodes in each sample were amplified using unique index primers. In total 23-31 PCR cycles (depending on the experiment and sample) were performed with 25 µL DNA (ChIP undiluted, input 1:50), 8 µL 5x High-Fidelity Phusion buffer (ThermoFisher), 0.4 µL dNTPs (10 mM), 0.4 µL Phusion High-Fidelity DNA Polymerase (ThermoFisher), 0.4 µL primer 1 (p5long-LTR2EF_U5, 10 µM), 0.4 µL primer 2 (sample specific index primer), 0.4 µL P1 Nextera (Illumina) primer (10 µM) and 0.4 µL P2 Nextera (Illumina) primer (10 µM) in a total volume of 40 µL. PCR products were quantified on a 2% agarose gel using ImageJ software. Subsequently, PCR products were mixed in equimolar fashion and purified from a 2% agarose gel with a QIAquick gel extraction kit (Qiagen). Purified DNA was sequenced (single read, 65 bp) on a HiSeq2500 platform (Illumina). For data analysis, samples were de-multiplexed using the 12 bp sample index resulting in 1.5-6M reads per sample. Barcode counting and data preprocessing and filtering were performed as described previously [8] with minor modifications. To optimize the normalization step and since most analyses only required a specific subset of the data, two subsets were created: 1) only tDNA samples (Epi-Decoder tDNA-Ty1 locus) named 'tDNA' and 2) only HO samples (Epi-Decoder HO locus) named 'ho'. In the 'tDNA' subset, counts of technical replicates were summed to a biological replicate, in the 'ho' subset the mean was taken. For normalization, the relative library size factor was calculated. For the 'tDNA' subset, this factor was based on the total library size and for the 'ho' subset on the sample median because the 'ho' subset is screened in sub pools. Additionally, a combined set of the two subsets was created: 'tDNA_ho'. The counts of the two underlying subsets were normalized using a relative size factor based on total library size. For differential analysis of enrichment (ChIP) in 'tDNA' and 'tDNA_ho', extra columns were created with ChIP (IP) values corrected for differences in input (IN). For this, a relative size factor was calculated per row (protein) for each of the IN conditions to correct the corresponding IP. Differential analysis was based on a paired analysis with DEseq2 [9] using the 'tDNA' subset with the following two exceptions. Differential analysis of IP and input at the HO locus in glucose was based on the 'ho'-subset and was done paired. Differential analysis of IP between tDNA and HO was done based on the 'tDNA_ho' subset using the corrected IP values and was done unpaired.

**Epi-Decoder validations**

Western blot and PCR were used to verify outliers and unexpected hits in the Epi-Decoder screens. We identified a few proteins that lacked a (functional) TAP-tag, were not clonal in the arrayed library, or had a wrong genetic background. **Supplemental Table S12** summarizes the excluded proteins. In addition, histone proteins were excluded from the analyses since they are present at very high levels, take up a major part of the read counts, and were spiked-in differently between experiments.

**ChIP-seq data analysis**

ChIP-sequencing reads were mapped to the sacCer3 reference genome downloaded from UCSC using the BWA-MEM program version 0.7.17-r1188 [10] using the option '-M'. No filtering for mapping quality was performed. Coverage tracks were created using the GenomicRanges [11] and rtracklayer

[12] R/Bioconductor packages. Specifically, unique reads were extended 200 bp from their 5' end and received a penalty for the number of alternative mapping positions. Coverage was down sampled to integrate to 1x genome size (reads per genomic content, RPGC). For counting reads in gene bodies, we took gene bodies from Ensembl's R64-1-1.108 GTF annotation [13] and extended these by 150 bp in both directions, specifically to more generously count reads at tDNA positions. Counting was then performed using the 'summarizeOverlaps' function from the GenomicAlignments R/Bioconductor package [11], wherein parameters were set such that the minimum mapping quality was greater than 10 and no duplicated reads were counted. To make the tornado heat map of ChIP-seq and ChIP-exo signal at tDNAs, excluding mitochondrial tDNAs, we used the tornadoplot R/GitHub package (version at hash ea8f4c6, https://github.com/teunbrand/tornadoplot) with arguments set to 'width = 2000, binwidth = 1' and sorted on Fpt1 signal. R version 4.2.1 and Bioconductor release 3.15 were used.

**Classification of housekeeping and regulated tRNA genes**

Based on UV cross-linking and analysis of cDNA (CRAC) to capture nascent RNAs bound by RNAPIII, two groups of tRNA genes have been identified by Turowski et al. (**Supplemental Table S6**) [2]. The CRAC dataset contains data for 275 tRNA genes. One canonical group of 'regulated tRNA genes' shows loss of engaged RNAPIII upon a switch from glucose to glycerol (RNAPIII CRAC gly/glu ratio < 1, here described as low RNAPIII retention score) and repression that is dependent on Maf1 (RNAPIII CRAC *maf1Δ*/wt ratio > 1, here described as high Maf1 dependency score). The other group of 'housekeeping tRNA genes' is defined as RNAPIII CRAC gly/glu ratio > 1 (high RNAPIII retention score), and RNAPIII CRAC *maf1Δ*/wt ratio < 1 (low Maf1 dependency score).

**ChIP-seq and ChIP-exo analysis**

Count data at individual tRNA genes from ChIP-seq and ChIP-exo and an overview of tRNAs included in the ChIP-seq heat maps can be found in **Supplemental Table S6**.

*ChIP-seq:* Genome tracks as in **Figure 2A-B and S2A** (bigwig files) were displayed in the UCSC genome browser. For Fpt1 counts at tRNA genes (as in **Figure 2I**) in the ChIP-seq dataset, tRNA genes with zero counts (n = 20) were excluded. In addition, tRNA genes that have gly/glu or *maf1Δ*/wt ratios classified as outliers (determined with a box plot) were not included (n = 12) resulting in 243 tRNA genes. The average Fpt1 occupancy as depicted in **Figure 2I** was determined based on the 243 included tRNA genes. The dataset used to calculate the Pearson correlation (as in **Figure S2D**) between gly/glu (RNAPIII retention score), *maf1Δ*/wt (Maf1 dependency score) and Fpt1 occupancy at tRNA genes was performed on this same subset of 243 tRNA genes.

*ChIP-exo*: Hypervariable tRNA genes were excluded from the analysis leaving 261 tRNA genes. Selection criteria and coordinate reference files were obtained from a previously published study to analyze Fpt1 (and related) datasets [14]. ChIP-exo plots were smoothened using 10 neighbors and 4$^{th}$ order (Prism 9 V9.4.1). Heat maps were based on NCIS normalized CDT files that contain strand-combined data. The dataset used to calculate the average Fpt1 occupancy as depicted in **Figure S2F** and the Pearson correlation (as in **Figure S2E**) between the gly/glu, *maf1Δ*/wt and Fpt1 occupancy at tRNA genes was filtered for outliers (determined with a box plot) in gly/glu and *maf1Δ*/wt ratios (n = 12) resulting in 249 tRNA genes.

**Live cell imaging microscope settings and data analysis**

Live cell imaging was performed on a setup consisting of an AxioObserver inverted microscope (Zeiss), an alpha Plan-Apochromat 100× NA 1.46 oil objective, an sCMOS ORCA Flash 4v3 (Hamamatsu) with a 475-570 nm dichroic (Chroma), 570 nm longpass beamsplitter (Chroma), 515/30 nm emission filter (Semrock for GFP) and 600/52 nm emission filter (Semrock for mScarlet), an UNO Top stage incubator (OKOlab) at 30°C, and LED excitation (SpectraX, Lumencor) at 470/24 nm at 100% power for 50 ms exposure (for GFP) and 550/15 nm at 100% power for 150 ms exposure time (for mScarlet-I) resulting in a 3.1 W/cm$^2$ excitation intensity for the GFP signal and 41.3 W/cm$^2$ for the mScarlet-I signal. For each position, a *z*-stack (9 slices, Δ*z* 0.5 µm) was recorded using Micro-Manager software [15]. For each condition, 3-5 images were taken for each of the 3 biological replicates, with in total at least 327 cells. The microscopy data was analyzed using custom-written Python software. First, a maximum intensity projection was made. Next, the mScarlet-I channel was used to segment cells using Otsu thresh-holding and water shedding. For each cell, the total nuclear mScarlet-I intensity was determined in each of the 9 slices in the *z*-stack, and the *z*-slice with the maximum signal was taken to be the slice where the nucleus is in focus. In this *z*-slice, the nuclear and cytoplasmic intensity were determined and background correction was performed on nuclear and cytoplasmic intensities by subtracting the median value of all intensities measured in the same *z*-slice outside the cells. For each cell, the nuclear enrichment was taken to be the ratio between the median nuclear and median cytoplasmic signal.

**Competitive growth assay and flow cytometry analysis**

Equally mixed NeonGreen and mScarlet labelled cells were inoculated in YNB + auxotrophic amino acids + 2% glucose (Glu-AA), SC + 2% ethanol (Glu/EtOH) and SC + 2% glycerol (Gly 37°C) and incubated at 30°C, except for glycerol which was incubated at 37°C. Glu-AA samples were switched to SC + 2% glucose every other day and Glu/EtOH to SC + 2% glucose for 24h after 48h of growth in SC + 2%

ethanol. Glycerol samples were diluted every 3-4 days to prevent saturation of the culture. At t = 0, 6 and 14 days, aliquots were collected to make glycerol stocks for storage at -80°C. To analyze samples with flow cytometry, samples were thawed and spun down for 3 minutes at 6000 rpm. Glycerol was removed and cells were resuspended in 1 mL SC + 2% glucose and briefly vortexed. Cells were sonicated for 10 seconds at level 3 (Diagenode, Bioruptor). DAPI was used as a dead-cell marker (1:50) and 50.000 events were measured for each sample on a BD LSRFortessa cell analyzer (BD Biosciences). The following laser-settings were used: BL(D) 530/30 (NeonGreen), YG(D) 610/20 (mScarlet) and V(F) 450/50 (DAPI). Data was processed and analyzed using FlowJo software. The Malthusian coefficient *m* was calculated as described in [16] using $m = \ln(10^{\log[(MUTend/WTend)/(MUTstart/WTstart)]/t})$ in which *MUTend* and *WTend* are the fraction of *fpt1Δ* and wild-type cells at t = 14, *MUTstart* and *WTstart* the fraction at t = 0 and *t* is the number of generations. The number of generations was determined based on known growth curves. The Malthusian coefficient was corrected for reporter effects determined by the wild-type/wild-type mixtures.

**Immunoblotting**

*Rpo31 immunoblot (8% polyacrylamide gel) – SUME prep*

Cells were grown in 15 mL YEPD to mid-log phase. Cell pellets were collected by a 5-minute spin at 2000 rpm and subsequently resuspended in media containing a different carbon source. Samples were harvested by spinning cells at 2000 rpm for 5 minutes. The pellet was washed with 1 mL TE + PMSF (1:500 100 mM). Pellets were stored at -80°C until further processing. Cell pellets were resuspended in 400 µL SUME buffer (1% SDS, 8M Urea, 10mM MOPS, pH 6.8, 10mM EDTA, 0.01% bromophenol blue) containing proteinase inhibitors (proteinase inhibitor cocktail, EDTA-free, Roche). Cells were lysed for 3 minutes by bead beating using zirconia silica beads (BioSpec, 0.5 mm). Samples were incubated for 10 minutes at 65°C and centrifuged at 13000 rpm for 10 minutes to pellet beads and insoluble proteins. Protein concentration of the supernatant was measured using a DC protein assay (Bio-Rad) according to the manufacturer's protocol. Approximately 20-30 µg protein was separated on an 8% polyacrylamide gel in 1x TGS (Tris-Glycine-SDS) running buffer. Proteins were transferred to 0.45 µM nitrocellulose blotting membrane for 4h using 1A at 4°C.

*Fpt1 immunoblot (10% polyacrylamide gel) – TCA prep*

Procedure was similar to Rpo31 immunoblot but with the following modifications. After pellet collection, the pellet was washed with 1 mL 20% TCA and resuspended in 200 µL 20% TCA (trichloroacetic acid). Cells were lysed for 3 minutes by bead beating using zirconia silica beads (Biospec, 0.5 mm). Beads were washed 3 times with 200 µL 5% TCA and supernatant was pooled. Cells were centrifuged for 10 minutes at 3000 rpm at 4°C. Pellets were resuspended in 100 µL Laemmli Buffer and 50 µL 2M Tris pH 8.5 was added to neutralize the solution. Samples were incubated for 5 minutes at 95°C and centrifuged for 10 minutes at 3000 rpm at RT. Protein extract (6 µL) was separated on a 10% polyacrylamide gel in 1x TGS (Tris-Glycine-SDS) running buffer. Proteins were transferred to 0.45 µM nitrocellulose blotting membrane for 2h using 1A at 4°C.

Membranes were blocked with 5% milk powder (Nutrilon) in PBS for 30 minutes at room temperature. Primary antibody staining for anti-TAP (Invitrogen CAB1001 rabbit anti-TAP, 1:2500) was done in 2% milk in TBS-T overnight at 4°C. After washing three times with TBS-T, membranes were incubated with the secondary antibody (goat anti-rabbit antibody IRDye 800, Licor, 1:10000) in 2% milk in TBS-T for 45 minutes. Pgk1 was stained with Pgk1 primary mouse antibody (Invitrogen 22C508, 1:2000) and secondary donkey anti-mouse antibody (IRDye 680, Licor, 1:10000) for 2h and 45 min respectively in 2% milk in TBS-T. Membranes were washed once in PBS before imaging on a LI-COR Odyssey IR Imager (LI-COR Biosciences). Quantification of gels was done using ImageJ software. Pgk1 protein levels were used to normalize Fpt1 and Rpo31 protein levels.

**Phylogenetic analysis**

Orthologous sequences of *YKR011C* were retrieved from 77 publicly available yeast genomes. For 41 genomes, gene model annotations and the corresponding amino acid sequences were publicly available ("annotated proteins" column in **Supplemental Table S13**). For the remaining 36 genomes, the protein annotations were not available, and putative orthologues of *YKR011C* were predicted with a sequence similarity search ("predicted protein" column in **Supplemental Table S13**). Putative orthologous were retrieved by stringent sequence similarity search (e-value < 10e-40) using EMBOSS (6.6.0) and BLAST (2.2.31+) and using the 24 previously identified *YKR011C* orthologues as query sequences. Yeast proteomes were clustered in gene families using OrthoFinder (2.2.7). Protein sequences were aligned with MAFFT (v7.271) and a Maximum Likelihood tree was inferred with IQ-Tree (1.6.8). *Hanseniaspora* sequences were highly divergent from other *YKR011C* orthologues and therefore dropped from the alignment.

**Statistical analyses**

Statistical analyses were performed using Prism 9 V9.4.1.

**Supplemental Table S9.** List of strains used in this study.

| Name | Genotype | Figure | Source |
|---|---|---|---|
| BY4741 | MATa his3Δ1 leu2Δ0 met15Δ0 ura3Δ0 | 3B, 4H, S4H, 6A-D, S6A-C | [17] |
| TAP-tag library | MATa his3Δ1 leu2Δ0 ura3Δ0 met15Δ0 yfp-TAP-His3MX6 | 1B-E, S1A-D, 5B-C, S5A-C | [18] |
| NKI4212 | MATα can1::HphMX lyp1::STE3pr-LEU2 his3Δ1 leu2Δ0 ura3Δ0 met15Δ0 | 1B-E, S1AB-D, 5B-C, S5A-C | [8] |
| NKI4214 (library) | MATα can1::HphMX lyp1::STE3pr-LEU2 his3Δ1 leu2Δ0 ura3Δ0 met15Δ0 yfp-TAP-His3MX6 | 1B-E, S1AB-D, 5B-C, S5A-C | [8] |
| NKI4217 (library) | MATα can1Δ::HphMX lyp1Δ::STE3pr-LEU2 his3Δ1 leu2Δ0 ura3Δ0 met15Δ0 yfp-TAP-His3MX6 ho::BC-KanMX-BC | 1C, S1A, S1C | [8] |
| NKI5601 | MATa his3Δ1 leu2Δ0 ura3Δ0 met15Δ0 chrXIII-192201::NatMX | 1B-E, S1B-D, 5B-C, S5A-C | This study |
| NKI5614 (library) | MATα can1::HphMX lyp1::STE3pr-LEU2 his3Δ1 leu2Δ0 ura3Δ0 met15Δ0 chrXIII-192201::NatMX tP(UGG)M::tP(UGG)M-barcode | 1B-E, S1B-D, 5B-C, S5A-C | This study |
| NKI5616 (library) | MATα can1::HphMX lyp1::STE3pr-LEU2 his3Δ1 leu2Δ0 ura3Δ0 met15Δ0 yfp-TAP-His3MX6 chrXIII-192201::NatMX tP(UGG)M::tP(UGG)M-barcode | 1B-E, S1B-D, 5C, S5C | This study |
| NKI5642 (library) | MATα can1::HphMX lyp1::STE3pr-LEU2 his3Δ1 leu2Δ0 ura3Δ0 met15Δ0 yfp-TAP-His3MX6 chrXIII-192201::NatMX tP(UGG)M::tP(UGG)M-barcode fpt1Δ::KanMX | 5B-C, S5A-C | This study |
| NKI5662-4 | MATα can1::HphMX lyp1::STE3pr-LEU2 his3Δ1 leu2Δ0 ura3Δ0 met15Δ0 RPO31-TAP-His3MX6 chrXIII-192201::NatMX tP(UGG)M::tP(UGG)M-barcode | 4A-I, S4A-H | This study |
| NKI5665-7 | MATα can1::HphMX lyp1::STE3pr-LEU2 his3Δ1 leu2Δ0 ura3Δ0 met15Δ0 RPO31-TAP-His3MX6 chrXIII-192201::NatMX tP(UGG)M::tP(UGG)M-barcode fpt1Δ::KanMX | 4C-I, S4B-H | This study |
| NKI5623 | MATα can1::HphMX lyp1::STE3pr-LEU2 his3Δ1 leu2Δ0 ura3Δ0 met15Δ0 FPT1-TAP-His3MX6 | 2A-I, S2A-G, 3A, S3A | This study |
| NKI4221 | MATα can1::HphMX lyp1::STE3pr-LEU2 his3Δ1 leu2Δ0 ura3Δ0 met15Δ0 RPL13A-TAP-His3MX6 | 2A-B, S2A | This study |
| NKI5625 | MATα can1::HphMX lyp1::STE3pr-LEU2 his3Δ1 leu2Δ0 ura3Δ0 met15Δ0 RPB2-TAP-His3MX6 | 2A-B, S2A | This study |
| NKI5626 | MATα can1::HphMX lyp1::STE3pr-LEU2 his3Δ1 leu2Δ0 ura3Δ0 met15Δ0 RPO31-TAP-His3MX6 | 2A-B, S2A-B | This study |
| NKI5632 | MATa his3Δ1 leu2Δ0 ura3Δ0 met15Δ0 FPT1-TAP-KanMX | 2C-H, S2C, S2E-G, 3A-C, S3A | This study |
| NKI5753 | MATa his3Δ1 leu2Δ0 ura3Δ0 met15Δ0 FPT1-yEGFP-KanMX | 3D, S3B-D | This study |
| NKI5756 | MATa his3Δ1 leu2Δ0 met15Δ0 FPT1-yEGFP-KanMX ura3Δ0::PP7-NLS-mScarlet-URA3 | 3D, S3B-D | This study |
| NKI5633 | MATa his3Δ1 leu2Δ0 ura3Δ0 met15Δ0 fpt1Δ::KanMX | 6A-D, S6C | This study |
| NKI5620 | MATa his3Δ1 leu2Δ0 ura3Δ0 met15Δ0 fpt1Δ::KanMX | 6A-D | Open Biosystems: YSC1053 |
| NKI2580 | MATa his3Δ1 leu2Δ0 ura3Δ0 met15Δ0 maf1Δ::NatMX | S6C | This study |

| | | | |
|---|---|---|---|
| NKI2581 | MATa his3Δ1 leu2Δ0 ura3Δ0 met15Δ0 fpt1Δ:: KanMX maf1Δ::NatMX | S6C | This study |
| NKI5707 | MATa his3Δ1 leu2Δ0 ura3Δ0 met15Δ0 X-2::TEF1p-ymNeonGreen-CYC1term | 6F, S6D | This study |
| NKI5712 | MATa his3Δ1 leu2Δ0 ura3Δ0 met15Δ0 X-2::TEF1p-ymScarlet-CYC1term | 6F, S6D | This study |
| NKI5724 | MATa his3Δ1 leu2Δ0 ura3Δ0 met15Δ0 fpt1Δ::KanMX X-2::TEF1p-ymNeonGreen-CYC1term | 6F, S6D | This study |
| NKI5726 | MATa his3Δ1 leu2Δ0 ura3Δ0 met15Δ0 fpt1Δ::KanMX X-2::TEF1p-ymScarlet-CYC1term | 6F, S6D | This study |

**Supplemental Table S10.** List of plasmids used in this study.

| Name | Description | Yeast marker | Source |
|---|---|---|---|
| pFvL029 | pFA6a-TAP-kanMX6 | KanMX | [19] |
| pFvL099 | natMX6 | NatMX | [20] |
| pML104 | Cas9-sgRNA-URA3 | URA3 | [21] Addgene #67638 |
| pML104-NatMX3 | Cas9-sgRNA-NatMX3 | NatMX | Kind gift from Patricia Wittkopp (Addgene #83477) |
| pET542 | yeNeonGreen | - | Kind gift from Evelina Tutucci |
| pET543 | ymScarletI | - | Kind gift from Evelina Tutucci |
| pRS400 | kanMX6 | KanMX | [22] |
| pKT127 | pFA6a-yeGFP-KanMX6 | KanMX | [23] Addgene #8728 |
| pTL306 | URA3-PP7-NLS-ymScarletI | URA3 | [24] |
| pIla011 | Cas9-tDNA_Ty1-URA3 | URA3 | This study |
| pMvB03 | Cas9-X_2-NatMX | NatMX | This study |

**Supplemental Table S11.** List of oligos used in this study.

| Name | Forward | Reverse | Notes |
|---|---|---|---|
| Fpt1-TAP | TAATAGCCATCATCCGCATAAAAATGGCAAAATATCATTTTCCATGGAAAAGAGAAG | GATGTAGATGCTATAAAAAGACATGTTTTCCCGGAAATAGGAATTCGAGCTCGTTTAAAC | Strain construction |
| Fpt1-GFP | TAATAGCCATCATCCGCATAAAAATGGCAAAATATCATTTGGTGACGGTGCTGGTTTA | GATGTAGATGCTATAAAAAGACATGTTTTCCCGGAAATAGTCGATGAATTCGAGCTCG | Strain construction |
| *fpt1Δ* | GAAAAAGAAGAAGAAATACCAAGTACAAAGAAAGGAAACCAGATTGTACTGAGAGTGCAC | GATGTAGATGCTATAAAAAGACATGTTTTCCCGGAAATAGCTGTGCGGTATTTCACACCG | Strain construction |
| *maf1Δ* | ATTATAGGTGTAAGACAAGGAAAATTCACAAATTAAAGTTTAAAACTGTGCGGTATTTCACACCG | CGCTCATTACTCCAAACGGATTTTTTTGCCTAAAGAATCACGACAAGATTGTACTGAGAGTGCAC | Strain construction |
| NeonGreen/ Scarlet X-2 locus | GCTGAAGATTTATCATACTATTCCTCCGCTCGTTTCTTTTTTCAGTGAGGTGTGTCGTGAGCGGATAACAATTTCACACAGGA | CCACCAAGACAATATCCTGAAAATCGAAATTCTCGCCAAGGCATTACCATCCCATGTAAGCGCCAGGGTTTTCCCAGTCACGAC | Strain construction |
| X-2 gRNA | GATCGGCGACTAGGAAGAGAGTAGGTTTTAGAGCTAG | CTAGCTCTAAAACCTACTCTCTTCCTAGTCGCC | Strain construction |
| ZOD1 | GCATTTTAAAAGCCGGGAAA | ATAAAGCCTCACCCCGAGTT | qPCR |

| | | | |
|---|---|---|---|
| SCR1 | GTGGGATGGGATACGTTGAG | TTTACGACGGAGGAAAGACG | qPCR |
| RPR1 | TTGTTCCGTTTGACTTGTCG | TGGAACAGCAGCAGTAATCG | qPCR |
| SNR6 | CGAAGTAACCCTTCGTGGAC | TCATCCTTATGCAGGGGAAC | qPCR |
| ETC2 | CATATCATTGGAGGCGGTGC | TGAGTCACATAGCGAGCTGG | qPCR |
| ETC6 | ACTCATCCAGGCTTTCTCGA | TCAGTATCACCCGGAAAGCT | qPCR |
| NoORF | GGCTGTCAGAATATGGGGCCGTAGTA | CACCCCGAAGCTGCTTTCACAATAC | qPCR |
| tP(UGG)M | GCCCAAAGCGAGAATCATAC | TCGGTGATTCCTTAGTGGCTA | qPCR |
| tR(CCG)L | GCTCCTCTAGTGCAATGGTT | GACTCGAACCCGGATCACAG | qPCR |
| tK(UUU)K | GGTGTAGTGTGATGGTGGTGA | TTCTTCGGAGGGCTAACTCA | qPCR |
| tT(CGU)K | GGCCAAAAGGGCTGAGTAAT | CAAATGGCACCTGCTTCATT | qPCR |
| tV(CAC)D | GGGGAGCACAATGAAACAAAGG | GGTGCAGCAAGTCTCAACTTTAC | qPCR |
| tP(UGG)A | GAGCCAAAAGCAATCTGACC | GACCACACGCCCGTACTAAT | qPCR |
| tM(CAU)E | GCGGACCTCAATATAAGCGA | CGCTTAGCCAACTTGAAAGAA | qPCR |
| tL(CAA)G1 | AGAACCGAAACATACAAATAAGTGGT | TGATCACAGAACCAAAAAGATAAAA | qPCR |
| NatMX Epi-Decoder locus | TCGGTCCAACTTTTTCACCTCTTCTCAACGCTGTTTCAGAAGATTGTACTGAGAGTGCAC | TTAGCCACTTATTCGAGTATATGGCGGAAATTAAATGTACCTGTGCGGTATTTCACACCG | Epi-Decoder |
| tDNA-Ty1 gRNA | GATCTTGAGTGCATAATGTTGAAAGTTTTAGAGCTAG | CTAGCTCTAAAACTTTCAACATTATGCACTCAA | Epi-Decoder |
| Amplify barcode oligo library | AAAGTTTTCTGACAACTACAATATT | TCCAACAACTAGTAACACGA | Epi-Decoder |
| Barcode oligo library | AAAGTTTTCTGACAACTACAATATTAAGCCCTTAACAACAATAATGAATATTGAGTGCATAATGTTGA[NNNNNNNNNNNNNNNN]barcodeAATGGAACTCTAAAATATCATTTGTTTAGTAGTATTCGTGTTACTAGTTGTTGGA | | Epi-Decoder |
| Sample specific index primer | CAAGCAGAAGACGGCATACGAGAT[XXXXXXXXXXXX]index[NNNNNNNNNNNN]UMIACAAATGATATTTTAGAGTTCCATT | | Epi-Decoder |
| p5long-LTR2EF_U5 | AATGATACGGCGACCACCGAGATCTACACGCCCTTAACAACAATAATGAATATTGAGTGC | | Epi-Decoder |
| Nextera P1 | AATGATACGGCGACCACCGA | | Epi-Decoder |
| Nextera P2 | CAAGCAGAAGACGGCATACGA | | Epi-Decoder |
| Custom sequencing primer | GCCCTTAACAACAATAATGAATATTGAGTGCATAATGTTGA | | Epi-Decoder |

**Supplemental Table S12.** Proteins excluded from Epi-Decoder analysis. For each protein, the corresponding figure in which the protein is excluded and reason of exclusion is indicated.

| Protein | Figure | Reason |
|---|---|---|
| Gea2 | 1B-E, S1A-D, 5B-C, S5A-C | Wrongly annotated |
| Ecm30 | 1B-E, S1A-D, 5B-C, S5A-C | Wrongly annotated |
| Rad9 | 1B-E, S1A-D, 5B-C, S5A-C | Wrongly annotated |
| Rpc53 | 1B-E, S1A-D, 5B-C, S5A-C | No TAP-tag |
| Tma10 | 1B-E, S1A-D, 5B-C, S5A-C | Wrong genetic background |
| Med8 | 1B-E, S1A-D, 5B-C, S5A-C | No TAP-tag |
| Rpl6A | 1B-E, S1A-D, 5B-C, S5A-C | No TAP-tag |

| | | |
|---|---|---|
| Ahp1 | 1B-E, S1A-D, 5B-C, S5A-C | No TAP-tag |
| Rap1 | 1B-E, S1A-D, 5B-C, S5A-C | No TAP-tag |
| Pdc5 | 1B-E, S1B-D, 5B-C, S5A-C | Heterogeneous population tDNA-Ty1 library |
| Rpc40 | 1C, S1A | No TAP-tag HO library |
| Tfc8 | 1C, S1A | No TAP-tag HO library |
| Tfc6 | 1C, S1A | No TAP-tag HO library |
| Rpb11 | 5B-C, S5A-C | No functional TAP-tag *fpt1Δ* |
| Spt6 | 5B-C, S5A-C | No TAP-tag *fpt1Δ* |

**Supplemental Table S13.** Genome versions of species described in the ML tree from Supplemental Figure S1E. Annotated proteins were retrieved from publicly available yeast genomes, predicted proteins were predicted using sequence similarity search as described in Supplementary Material & Methods.

| Species | Version | Annotated proteins | Predicted proteins |
|---|---|---|---|
| *Candida glabrata* | ASM254v2 | X | |
| *Candida bracarensis* | GCA_900496985.1 | | X |
| *Nakaseomyces nivariensis* | ASM104691v1 | | X |
| *Lachancea fermentati* | GCA_900074765.1 | X | |
| *Lachancea thermotolerans* | ASM14280v1 | X | |
| *Eremothecium sinecaudum* | CP014249.1 | | X |
| *Eremothecium gossypii* | ASM9102v4 | X | |
| *Eremothecium coryli* | AZAH01.1 | | X |
| *Eremothecium cymbalariae* | ASM23536v1 | X | |
| *Kluyveromyces aestuarii* | AEAS01.1 | | X |
| *Kluyveromyces nonfermentans* | ASM367015v1 | | X |
| *Kluyveromyces wickerhamii* | AEAV01.1 | | X |
| *Kluyveromyces marxianus* | GCF_001417885.1 | X | |
| *Kluyveromyces dobzhanskii* | CCBQ01.1 | | X |
| *Kluyveromyces lactis* | ASM251v1 | X | |
| *Zygotorulaspora florentina* | ASM367157v1 | | X |
| *Zygotorulaspora mrakii* | ASM367156v1 | | X |
| *Torulaspora microellipsoides* | GCA_900186055.1 | | X |
| *Torulaspora delbrueckii* | ASM24337v1 | X | |
| *Torulaspora franciscae* | GCA_003705175.2 | | |
| *Torulaspora pretoriensis* | PPKE01.1 | | X |
| *Naumovozyma castelli* | ASM23734v1 | X | |
| *Naumovozyma dairenensis* | ASM22711v2 | X | |
| *Kazachstania africana* | GCF_000304475.1 | X | |
| *Tetrapisispora phaffii* | ASM23690v1 | X | |
| *Vanderwaltozyma polyspora* | ASM15003v1 | X | |
| *Kazachstania saulgeensis* | ASM90018042v1 | X | |
| *Kazachstania naganishii* | ASM34898v1 | X | |
| *Kazachstania unispora* | ASM370852v1 | | X |
| *Saccharomyces servazzii* | ASM221493v1 | | X |
| *Saccharomyces pastorianus_v1* | GCA_001515485.2 | | X |
| *Saccharomyces eubayanus* | GCF_001298625.1_SEUB3.0 | X | |
| *Saccharomyces bayanus* | ASM16703v1 | X | |

| | | | |
|---|---|---|---|
| *Saccharomyces uvarum_v1* | GCA_002242645.1 | X | |
| *Saccharomyces uvarum_v2* | ASM224264v1 | X | |
| *Saccharomyces kudriavzevii* | gca_000167075.IFO1802_v1.0 | X | |
| *Saccharomyces arboricola* | gca_000292725.SacArb1.0 | X | |
| *Saccharomyces mikatae* | SRX055454 | X | |
| *Saccharomyces paradoxus* | ASM207914v1 | | X |
| *Saccharomyces pastorianus_v2* | GCA_001515485.2 | | X |
| *Saccharomyces cerevisiae_v1* | GCA_000146045.2 | X | |
| *Saccharomyces cerevisiae_v2* | https://www.yeastgenome.org/reference/S000216438 | X | |
